# Reduced *SH3RF3* may protect against Alzheimer’s disease by lowering microglial pro-inflammatory responses via modulation of JNK and NFkB signaling

**DOI:** 10.1101/2024.06.23.600281

**Authors:** Ronak Patel, Rong Cheng, Christopher L. Cardona, Ellen Angeles, Gunjandeep Singh, Sabrina Miller, Archana Ashok, Andrew F. Teich, Angel Piriz, Aleyda Maldonado, Ivonne Jiménez-Velázquez, Richard Mayeux, Joseph H. Lee, Andrew A. Sproul

## Abstract

Understanding how high-risk individuals are protected from Alzheimer’s disease (AD) may illuminate potential therapeutic targets. We identified protective genetic variants in *SH3RF3*/*POSH2* that delayed the onset of AD among individuals carrying the *PSEN1*^G206A^ mutation. *SH3RF3* acts as a JNK pathway scaffold and activates NFκB signaling. While effects of *SH3RF3* knockdown in human neurons were subtle, including decreased ptau S422, knockdown in human microglia significantly reduced inflammatory cytokines in response to either a viral mimic or oAβ42. This was associated with reduced activation of JNK and NFκB pathways in response to these stimuli. Pharmacological inhibition of JNK or NFκB signaling phenocopied *SH3RF3* knockdown. We also found *PSEN1*^G206A^ microglia had reduced inflammatory response to oAβ42. Thus, further reduction of microglial inflammatory responses in *PSEN1*^G206A^ mutant carriers by protective variants in *SH3RF3* might reduce the link between amyloid and neuroinflammation to subsequently delay the onset of AD.

**Graphical Abstract:** 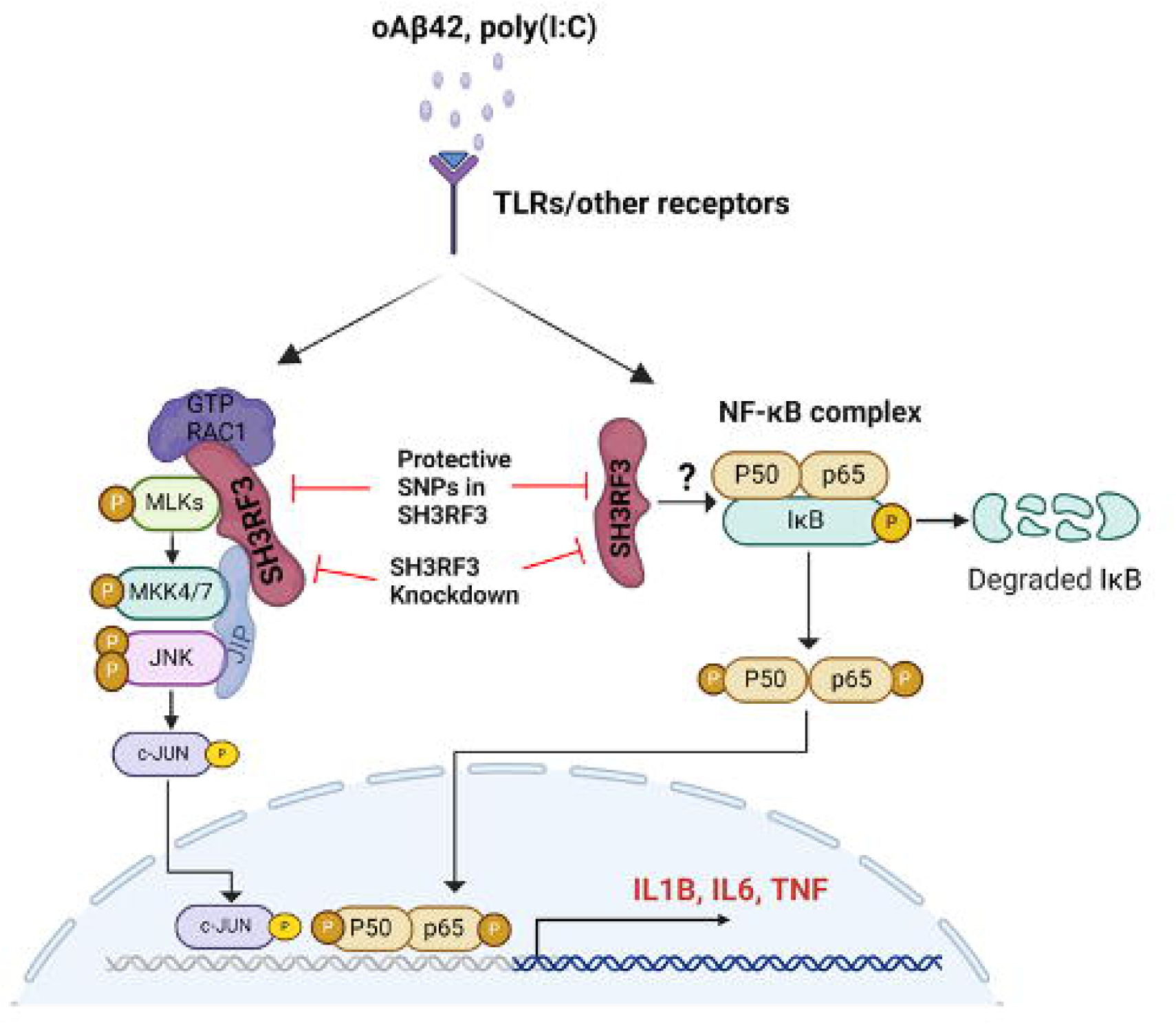

## Introduction

The amyloid hypothesis^1^ has largely guided therapeutic approaches to treat Alzheimer’s disease (AD) the past few decades. Although use of monoclonal antibodies to target Aβ removal has recently gained traction and FDA approval, clinical benefits are modest and have risks such as ARIA-E/H (amyloid-related imaging abnormalities-edema/hemorrhage)^2,3^. Thus, additional therapeutic strategies are necessary. Instead of targeting neuropathology directly, an alternative tactic is to better understand why some individuals are resilient to AD and then attempt to mimic this resistance via pharmacological manipulation or other interventions. One robust approach to understand resilience is the study of genetic factors which modify the age at onset of AD in high-risk populations such as presenilin-1 (*PSEN1*) mutation carriers, which is sufficient to cause early-onset AD (EOAD).

To date, much of the genetic work has been focusing on identifying risk variants, and a limited number of studies have made efforts to identify protective genetic factors^4,5^. As commented by Schwartz and colleagues^6^, protective genetic modifiers are likely to provide important insights into our understanding of the disease mechanisms of AD in the era of precision medicine. Our previous examination of phenotypic variation in the families with the G206A mutation in the *PSEN1* gene revealed that carriers of this mutation may display age at onset ranging from 40s to 80s even within the same family, and environmental risk factors explained little of the phenotypic variance. Our linkage analysis localized 2 loci, 2q13 and 4q35, that were likely to be associated with variation in age at onset of AD. Further examination of 2q13 identified *SH3RF3*/*POSH2* (*<u>p</u>lenty <u>o</u>f <u>SH</u>3 domains 2*) as one of the potential genetic modifiers that ameliorate the effects of the G206A mutation in *PSEN1*^4^. Among *PSEN1*^G206A^ carriers, *SH3RF3* was associated with a delay in age at onset of symptoms by 9.3 years. We further showed that variants in *SH3RF3* were associated with 0.8 to 2.9 years in delay in age at onset of symptoms in late onset AD (LOAD). This suggests that *SH3RF3* may serve as a genetic modifier in both early-onset AD (EOAD) and late-onset AD (LOAD), but the protective effect, represented by the mean years of accelerated or delayed onset is likely to be smaller in LOAD when compared with EOAD. Along with *SH3RF3*, we identified six other candidate genetic modifiers in 2q13 and 4q35 that were nominally significant in both EOAD and LOAD, suggesting that modifiers that were discovered in EOAD may be generalizable to LOAD, the common form of AD.

We have focused this study on better understanding of the mechanisms of how *SH3RF3* may protect against AD. In addition to our study showing effects on AD risk^4^, *SH3RF3* has also been identified by causal inference analysis as a key microglial-expressed regulator of late-onset AD (LOAD)^7^ and in a genome-wide association study (GWAS) focused on HIV-associated neurocognitive disorder (HAND)^8^. SH3RF3 functional domains include 4 protein-protein interaction SH3 domains, a novel Rac1 binding domain that only binds the active GTP-bound version, and a RING domain which give SH3RF3 E3 ligase activity to target itself (and potentially other proteins) for proteasomal degradation. There are 3 *SH3RF*/*POSH* family members, which also include *SH3RF1*/*POSH* which has the same domain structure as *SH3RF3*, and *SH3RF2*/*POSHER* (POSH-eliminating RING *protein*), which is missing the 4^th^ SH3 domain. Both SH3RF1 and SH3RF3 have been identified as scaffold proteins promoting JNK (c-Jun N-terminal kinase) signaling as well as having the ability to activate NFκB signaling by an unknown mechanism when overexpressed^9–11^. The ability to act as a JNK scaffold has best been described for SH3RF1^12,13^, which directly binds GTP-Rac1 and MAP3Ks such as MLK3 as well as JIP (*JNK-interacting protein*) family member scaffolds. JIP scaffolds, in turn, directly bind MAP2Ks (MKK4/7) and JNKs, which then phosphorylate c-Jun to activate the AP-1 transcription factor dimer (see also Graphical Abstract). On the other hand, SH3RF2/POSHER was reported to target SH3RF1/POSH for proteasomal degradation and interfere with its ability to promote JNK signaling^14^.

JNK and NFκB signaling pathways have both been shown to be strongly associated with AD pathogenesis and are activated by Aβ and upregulated in AD brains^15–18^. In neurons, JNK signaling has been shown to directly phosphorylate APP at T688 to promote amyloidogenic processing^19–21^ and tau at multiple residues^22,23^, increase BACE1 transcription^24^, and is required for cell death of murine neurons from oligomeric Aβ42^25^. In microglia, JNK signaling has been reported to mediate Aβ-driven cytokine production^26^, and has its activity increased by reduced *TREM2* expression^27,28^. Interestingly, NFκB signaling has also been reported to drive *BACE1* transcription and thus amyloidogenic processing^29^. The major pathogenic role of NFκB is that of a key driver of inflammatory responses such as the production of pro-inflammatory cytokines and reactive oxygen species (ROS) in glial cells, which is both detrimental in itself to neurons and promotes amyloid and tau pathology as well as damage to the blood brain barrier^17,18^. Importantly, a recent study using human induced pluripotent stem cell (hiPSC)-derived microglia (iMGLs) xenotransplanted into an AD mouse model (APP^NL-G-F^) demonstrated that human microglia have unique cytokine response modules (CRMs) in response to amyloid not seen in murine microglia^30^. CRMs are distinct from the DAM response by psuedotime analysis, are associated with NFκB signaling and pro-inflammatory cytokine production and include SH3RF3 as a member (CRM2). Therefore, we postulated that reduced *SH3RF3* expression or function would be protective against AD by mitigating extended activation of both JNK and NFκB pathways and tested this hypothesis in both hiPSC-derived neuronal and microglial (iMGLs) models.

## Results

### Identification of additional AD protective SNPs in *SH3RF3*

To further validate our initial discovery SNP in *SH3RF3*^4^, we significantly expanded our findings from the discovery *PSEN1*^G206A^ families to additional cohorts to validate and to generalize to the more common forms of AD.

#### Study Participants: Discovery dataset

The present study examined 505 study participants from 78 families from Puerto Rican with at least one carrier with the G206A mutation in the *PSEN1* gene (**Table 1**). The study design and description of the families were provided in detail in our earlier work^4^. Briefly, the mean age at onset of AD was 60.8 (SD=10.6), but the age at onset ranged from 40 to 93, depending on the carrier status. In these families, 50.9% of the family members carried the *PSEN1*^G206A^ mutation, and 40.6% of the family members were affected with AD where the mean age at onset was 58.1(8.1) for *PSEN1*^G206A^ carriers and 70.5(13.0) for *PSEN1*^G206A^ non-carriers. The *APOE4* allele frequency was 17.7 %. The mean years of education for these family members were 11.8 years (SD=4.9). We performed whole genome sequencing on approximately two-thirds (n=337) of the family members, and the remaining one-third (n=168) had the genome data generated from microarray SNP data with TOPmed imputation. As shown in **Table 1**, mutation carriers were more likely to be subjected to whole genome sequencing than non-carriers.

**Table 1.**
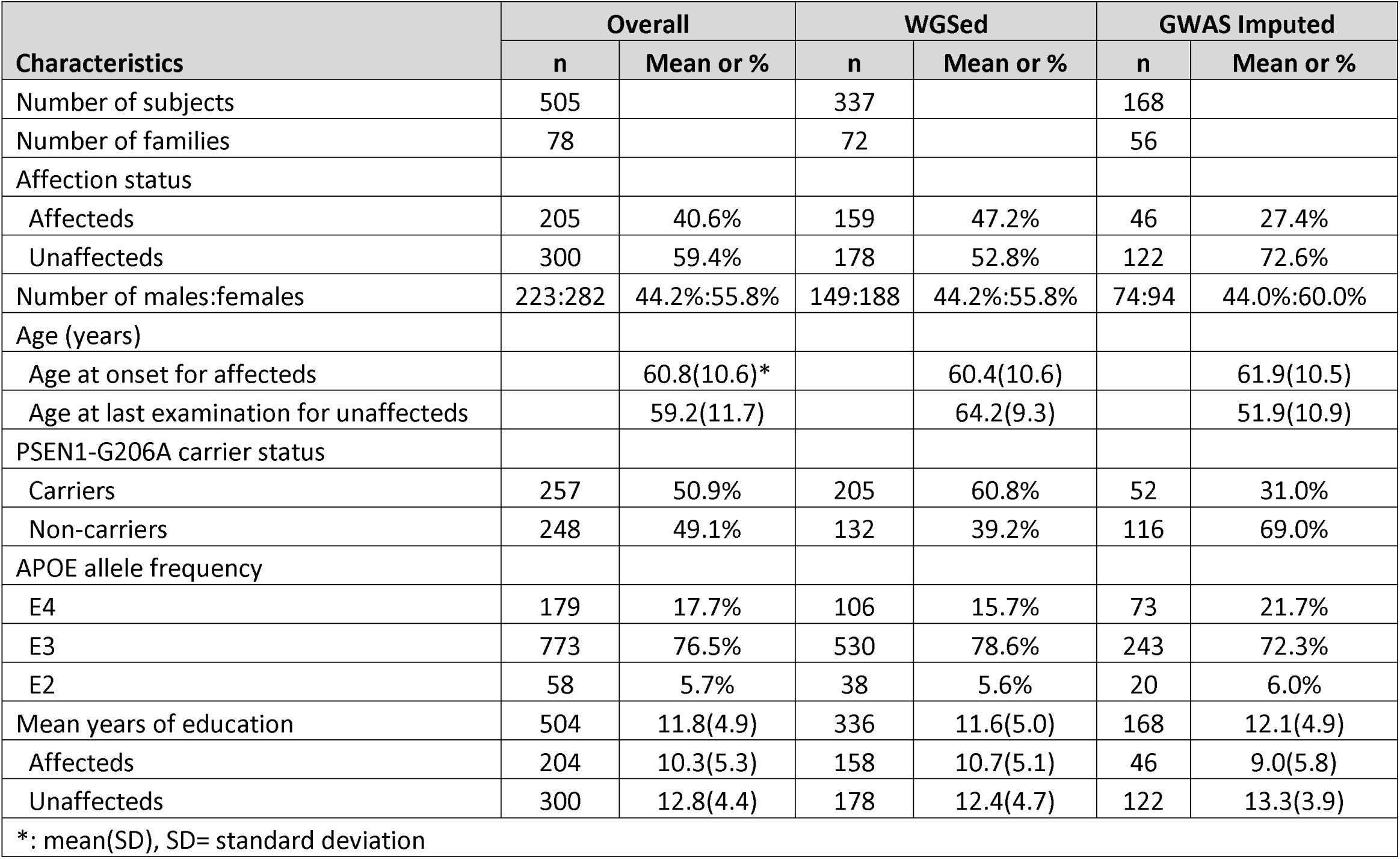
Demographic and clinical characteristics of the PSEN1 family study participants.

#### Allelic Association

Broadly, our multivariable allelic association model revealed that a tightly linked chromosomal region starting from 109,128,058 bp to 109,486,830 bp, encompassing 350kb were significantly associated with delay in age at onset of AD as shown in **Table 2**. Furthermore, when stratified by the *PSEN1*^G206A^ status, three contiguous sets of SNPs were associated with delay in age at onset of AD, with medians ranging from 5.4 years to 18.6 years. In contrast, no significant delay in age at onset was observed in non-carriers of the *PSEN1*^G206A^ mutation. This striking difference in the observed delay in age at onset supports a possible interaction between *PSEN1* and *SH3RF3*. Specifically, a long stretch region was associated with approximately 5-year delay in age at onset of AD from chromosome location 109,288,547 to 109,387,052 bp in carriers (region 2), and the low allele frequency suggested that a few haplotypes may be segregating in a limited number of families. Two other regions indicated in green -- 109,128,058 to 109,263,859 (region 1) and 109,399,707 to 109,486,830 (region 3) – showed significant association with stronger effect size; however, the variability was greater than that in region 2. We, however, note that regions 1 and 3 contain synonymous mutation at 109,130,101 bp (rs77039993) and nonsynonymous mutation at 109,449,440 bp (rs192679474) with significantly larger effect size (18.6 and 25.4, respectively). Three individuals had a synonymous variant at 109,130,101bp, and one had a nonsynonymous variant at 109,449,440bp.

**Table 2.**
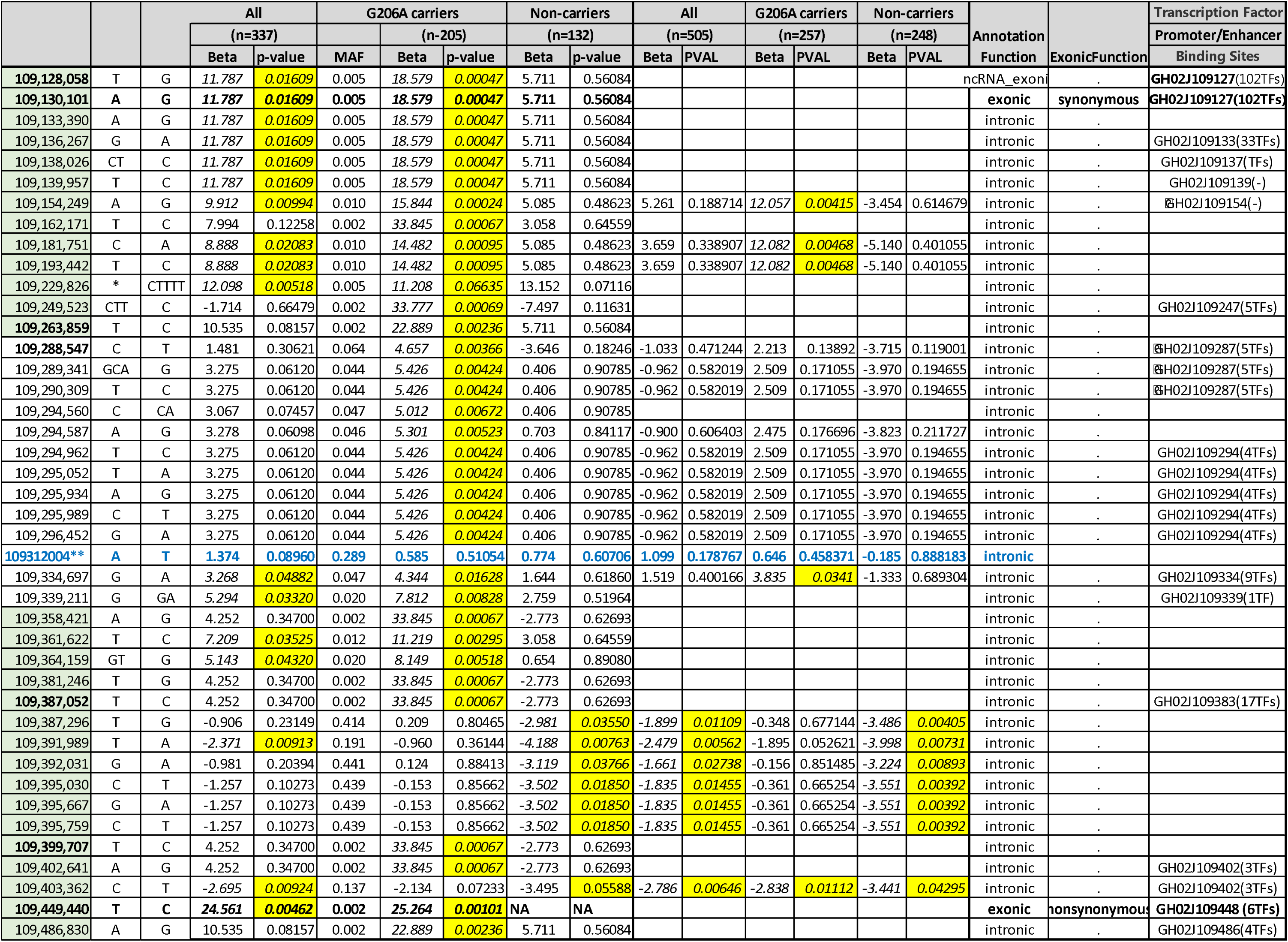
Allelic Association with Age at Onset or AD or age at last follow up for SH3RF3: The whole genome sequencing data and combined with GWAS with imputation.

#### Haplotype in gene enhancer regions

Given that these three regions represent primarily noncoding regions, we further examined each variant to see functional relevance (**Table 2**). Of 15 SNPs that were significantly associated, more than one-half (12) were located in enhancer regions.

#### Replication Datasets: Familial LOAD and Sporadic AD

We then examined two other datasets, namely two cohorts of Estudio Familiar de Influencia Genetica en Alzheimer (EFIGA)^31^ and late onset sporadic AD (WHICAP^32^) to determine whether the observed delay in age at onset can be confirmed in independent datasets, and then be generalized to individuals without an autosomal dominant mutation, *PSEN1*^G206A^. Since these familial LOAD and sporadic AD cases do not carry the *PSEN1*^G206A^ mutation, we used *APOE4* as the risk variant instead. In these two lower risk cohorts (**Figure 1; Table S1)**, SNPs in SH3RF3 were consistently associated with delay in age at onset in *APOE4 carriers,* but the magnitude of delay in age at onset was smaller (range: 1.3-2.9 years) than that in the *PSEN1*^G206A^ carriers. As in the *PSEN1*^G206A^ carrier families, allelic associations were not significant in non-carriers of *APOE4*. This parallels our earlier work in *PSEN1*^G206A^ mutation carriers, suggesting that genetic modifiers identified from *PSEN1*^G206A^ mutation carriers can be generalized to other forms of AD. We further note that these variants are located within the gene enhancer regions, suggesting that they may influence the gene expression.

**FIgure 1:**
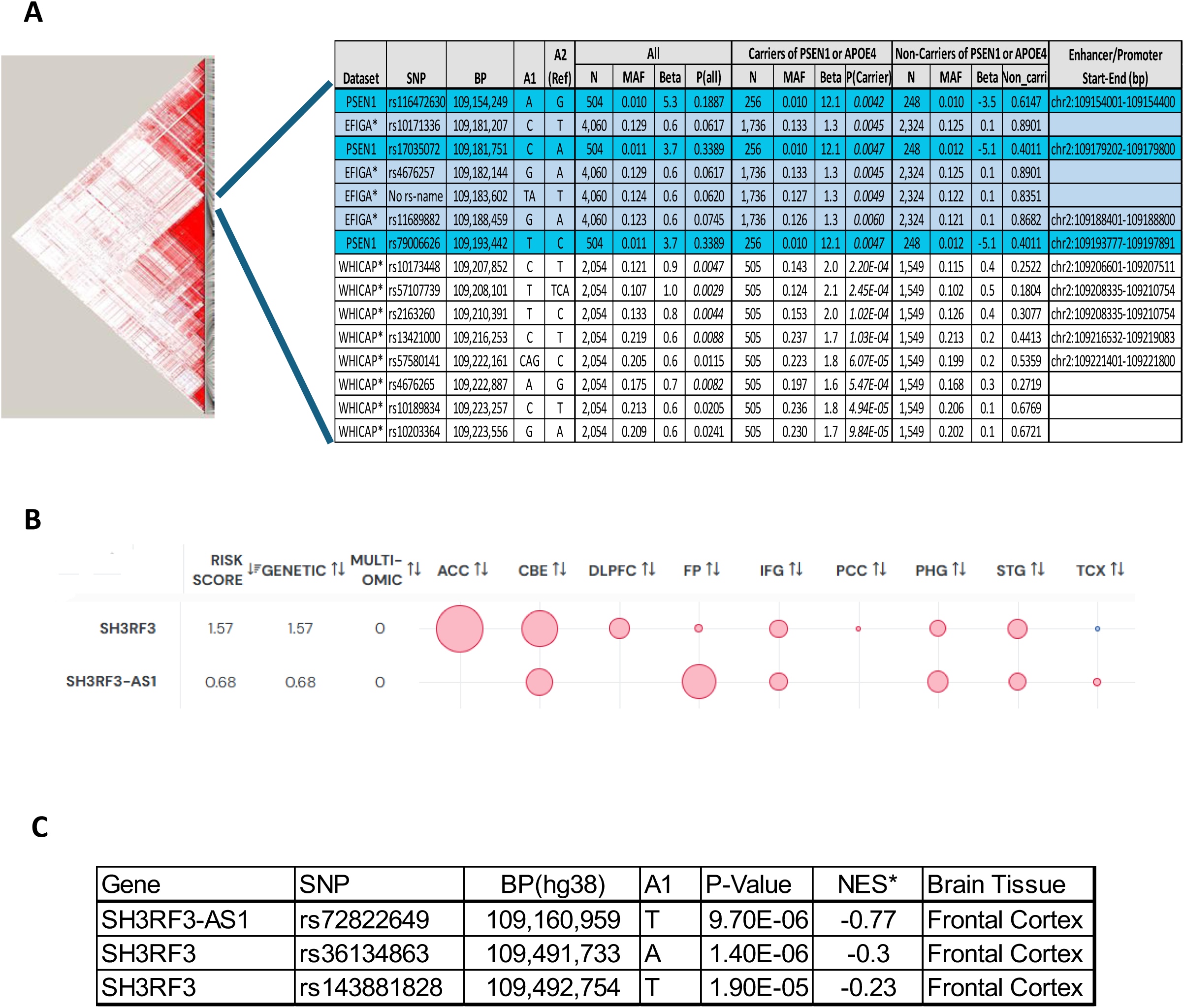
Comparisons of genetic association in *SH3RF3* by cohorts PSEN1G206A, EFIGA, and WHICAP, and differential gene expression patterns. (A) Identification of SNPs associated with delay in age at onset of AD in *APOE4* carriers in EFIGA and WHICAP studies. (B) *SH3RF3* is down-regulated in anterior cingulate cortex and cerebellum, and *SH3RF3*-AS1 is down regulated in frontal pole. Data is normalized for age of death and for circles, red indicates reduced expression and the diameter reflects p-value. Anterior Cingulate Cortex (ACC), Cerebellum (CBE) Dorsolateral Prefrontal Cortex (DLPFC), Frontal Pole (FP), Inferior Frontal Gyrus (IFG), Posterior Cingulate Cortex (PCC), Parahippocampal Gyrus (PHG), Superior Temporal Gyrus (STG), and Temporal Cortex (TCX). (C) Three SNPs in the frontal cortex were significantly downregulated in GTEx.

#### Differential Gene Expression in LOAD

We used the AGORA website, a publicly available transcriptomic, proteomic, and metabolomic dataset that examines its association with AD in brain tissues (https://agora.adknowledgeportal.org). As shown in **Figure 1B**, the log2FCs for the *SH3RF3* gene and the adjacent *SH3RF3*-AS1 gene was downregulated in all tissues; however, only three tissues, namely Anterior Cingulate Cortex (ACC) and cerebellum (CBE) had adjusted p_adjusted_ <0.05 for *SH3RF3.* In addition, frontal pole was similarly downregulated for *SH3RF3*-AS1. The SNPs in *SH3RF3* and *SH3RF3-AS1* were downregulated in certain brain tissues of affected individuals when compared with those of unaffected. Nearly one-half of the SNPs in bold face in **Table 2** were in promotor or enhancer regions. We then further examined SNPs in regions 1-3 in GTEx and identified three SNPs that were differentially expressed by genotype in frontal cortex (**Figure 1C**). It is equally important to note that those three SNPs were not differentially expressed in other brain tissues. This suggests the effects of *SH3RF3* on AD pathogenesis are not primarily driven by transcriptional responses to AD pathology globally but rather by regulation of SH3RF3 protein function or levels. In addition, the role of *SH3RF3*-AS1 (anti-sense) is unclear and should be the focus of future studies.

### *SH3RF3* knockdown does not affect APP processing in human neurons

We next sought to understand the mechanisms of how protective SNPs in *SH3RF3* affected AD risk using human model systems. We first assessed loss of *SH3RF3* function in neurons produced by NGN2-mediated transdifferentiation of hPSCs (H9 hESC line) into excitatory neurons using a permanent line method developed by our lab ^33^. We used pooled siRNAs to knockdown (KD) *SH3RF3* in neurons which was verified by qPCR (**Figure 2A**). We then tested for potential JNK pathway-related modulation of APP and Aβ production that might be modulated by *SH3RF3* KD. In addition to the effects on APP processing described above for JNK signaling, APOE isoforms can differentially activate the AP1 transcription factor to increase APP transcription and Aβ production^34^. The downstream JNK target c-Jun can also activate AP1 when phosphorylated and dimerized with FOS. *SH3RF3* KD resulted only in a trend of reduced APP transcription and total Aβ production (**Figure 2B-C**). Importantly, the Aβ42:Aβ40 ratio also remained unchanged (**Figure 2D**), and there was no effect on pAPP T688 (**Figure S2A**). This suggests that while protective SNPs in *SH3RF3* may have subtle beneficial effects on modulating APP transcript levels, overall, it is unlikely that their protection against AD is primarily mediated through effects on Aβ production.

**Figure 2:**
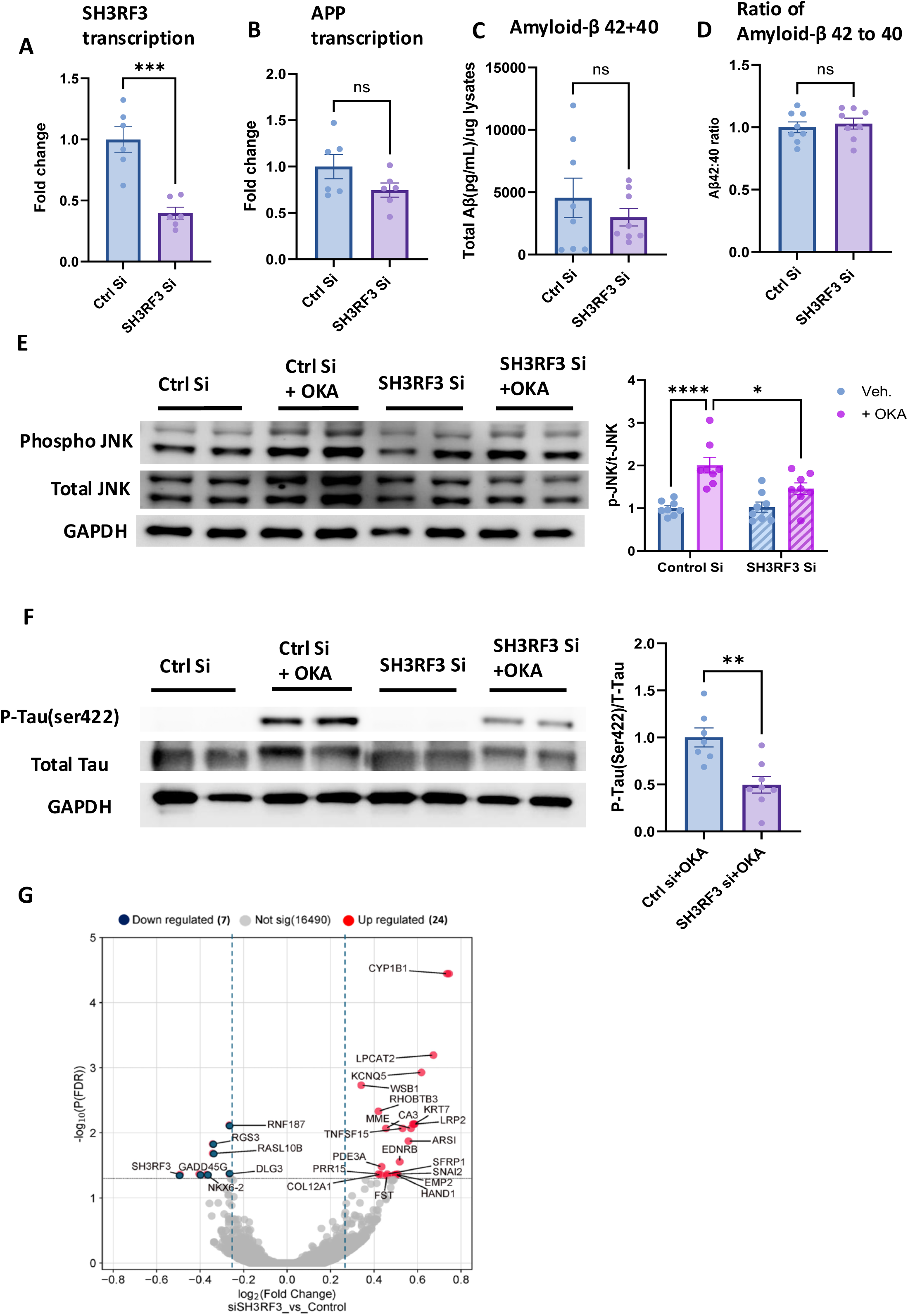
*SH3RF3* KD does not affect APP processing but does reduce pTau S422. (A-B) qPCR analysis of *SH3RF3* and APP in hPSC-derived neurons treated with non-targeting control or *SH3RF3* siRNA. *SH3RF3* is significantly knocked down but subsequent effects on APP are only a trend. Data is presented as mean±SEM with N=3 biological replicates (independent treated wells) for two independent experiments. Comparisons are made with unpaired t-test. (C-D) Multiplex ELISA for Aβ from conditioned media (CM) collected from neurons treated with either *SH3RF3* or control siRNA. Total Aβ (Aβ42+ Aβ40) levels for are similar for each condition. (C), as is Aβ42/Aβ40 ratio (D). Data is presented as mean±SEM with N=2-3 biological replicates per experiment for three independent experiments (N=8). Comparisons are made withunpaired t-test. (E-F) Western blot analysis of neurons treated with *SH3RF3* or control siRNA subjected to okadaic acid (OKA) or vehicle for 3h. Representative blots and quantification is shown. JNK signaling is strongly activated by OKA as measured by pJNK/JNK ratio (E), which is significantly reduced by *SH3RF3* KD. Data is presented as mean±SEM with N=2-3 biological replicates per experiment for three independent experiments (N=8), Comparison of control Si with OKA and *SH3RF3* Si with OKA by two-way ANOVA, ****<0.0001, *<0.05. pTau S422 phosphorylation (relative to total tau) is strongly induced by OKA treatment (F), which is decreased by *SH3RF3* KD. Data is presented as mean±SEM with N=2 or 3 biological replicates per experiment from 3 independent experiments (N=7-8). Conditions without OKA have undetectable bands for pTau S422 and only conditions with OKA are quantified by comparison of control Si with OKA and *SH3RF3* Si with OKA by unpaired t-test, **<0.01. (G) Heatmap showing differently expressed genes upon *SH3RF3* knockdown in hPSC-derived neurons, pAdj<0.05

### *SH3RF3* knockdown decreases JNK-specific tau phosphorylation at ser422

We then assessed for effects of *SH3RF3* KD on tau phosphorylation. We first confirmed *SH3RF3* KD modulates JNK pathway activity in human neurons under basal and JNK pathway-inducing conditions. The protein phosphatase 2A inhibitor okadaic acid (OKA) activates JNK signaling in cultured hippocampal neurons and induces hyperphosphorylated tau and neuronal cell death^35,36^. OKA induced significant activation of JNK signaling in human neurons, which was moderately reduced by *SH3RF3* KD (**Figure 2E**). S422 has been identified as a prominent JNK-induced phosphorylation site in tau by mass spectrometry^23^. Tau S422 is expressed at very low levels under physiological conditions and co-localizes with PHFs in AD brains^37^, and correlates with tangle formation in murine models of frontotemporal dementia (FTD) and increased tau aggregation^38,39^. While pTau S422 was not detectable under basal conditions, *SH3RF3* KD under OKA treatment conditions demonstrated a reduction in pTau s422 (**Figure 1F**). On the other hand, there was only a non-significant trend of decreased pTau T217, a potential blood marker of AD^40,41^, from *SH3RF3* KD under OKA treatment conditions (**Figure S2B**). This suggests that protective SNPs that reduce *SH3RF3* expression levels or function may modulate AD risk in part by affecting levels of phosphorylation levels of tau at specific phospho sites.

### Transcriptomic analysis of *SH3RF3* KD in human neurons

We further assessed global gene expression profile of control siRNA and *SH3RF3* siRNA treated human neurons to identify other potential molecular phenotypes. We identified 31 genes with significant differential expression (p_Adjusted_<0.05) from *SH3RF3* KD (**Figure 2G**), which included eight genes previously associated with genetic effects on AD risk (unpublished data). Associated GO terms using DAVID included ubiquitin protein ligase activity and metal ion binding ^42^. Overall, this suggests that transcriptional effects from reduced *SH3RF3* expression in neurons might have small effects on AD risk, although this is most likely not the primary mechanism of protective SNPs.

### *SH3RF3* KD reduces inflammatory cytokines produced by iMGLs treated with the viral mimic poly(I:C) by modulating JNK and NFκB signaling

We then tested if *SH3RF3* KD had a more robust effect in microglia, where JNK and NFκB signaling have critical roles under pro-inflammatory conditions^43,44^. Further support from this approach came from looking at interactive published data sets of human and mouse CNS cell types, which showed that in the human brain *SH3RF3* had the highest expression in microglia and fetal astrocytes (**Figure S3A;** brainrnaseq.org)^45^. Interestingly, this was not the case in the mouse brain, where expression was highest in neurons and low in microglia (**Figure S3B**)^46^, suggesting species-specific differences in *SH3RF3*’s functional role.

Multiple neurodegenerative diseases have shown strong associations with viral exposure. This includes AD, which is correlated with viral encephalitis, herpes simplex infections, HHV6, picornaviruses and HIV type 1^47,48^. Viral infections are postulated to increase Aβ production as a protective response leading to increased amyloid deposition and induction of inflammatory molecular pathways in the CNS^49^. This idea is supported by recent findings that show increase of Aβ in brains from subjects infected by COVID-19^50,51^. Various Pathogen Associated Molecular Patterns (PAMPs) can activate Toll-Like Receptors (TLRs) on the surface of microglia leading to microglial activation and release of inflammatory cytokines in response to microbes likes viruses and/or Aβ accumulation, which in turn cause neurotoxic damage^52^. Increased release of pro-inflammatory cytokines is also associated with damage to oligodendrocytes and BBB leakage, leading to exacerbation of AD pathology^53,54^. Inhibitors blocking microglial activation and release of inflammatory cytokines have been proposed as a potential AD therapeutic strategy^55^.

Thus, we utilized a viral mimic poly(I:C) to treat iMGLs, double-stranded RNA (dsRNA) known to elicit inflammatory microglial response via TLR3 activation^56^. iMGLs were differentiated from the FA0000010 (FA10) hiPSC line using a previously published protocol with minor modifications^57^. Most differentiated cells were CD45^+^/CD11B^+^ as measured by flow cytometry (**Figure S3C**) and expressed key microglial markers such as Iba1 and P2RY12 by immunostaining (**Figure S3D**). We used similar pooled non-targeting control siRNA and siRNA targeting *SH3RF3* in iMGLs to knockdown *SH3RF3* in presence and absence of poly(I:C) stimulation. We observed moderate knockdown of *SH3RF3* by qPCR (**Figure 3A**). For assessing effects on pro-inflammatory cytokine production, we focused on 3 key Inflammatory cytokines, IL1β, IL6 and TNFα, which are also members of the human-specific CRM2 microglial cluster that appears in response to amyloid^30^. IL1β, IL6 and TNFα transcript levels upon 2h poly(I:C) stimulation showed strong upregulation compared to vehicle treated iMGLs, which were significantly reduced by *SH3RF3* KD in poly(I:C) conditions (**Figure 3B-D, Figure S3E).** This also translated to the protein level as measured by multiplex ELISA for IL1β, IL6 and TNFα from conditioned media from iMGLs at 16h post-treatment. Poly(I:C) treatment resulted in significantly higher release of IL1β, IL6 and TNFα which was significantly reduced by *SH3RF3* KD (**Figure 3E-G).**

**Figure 3:**
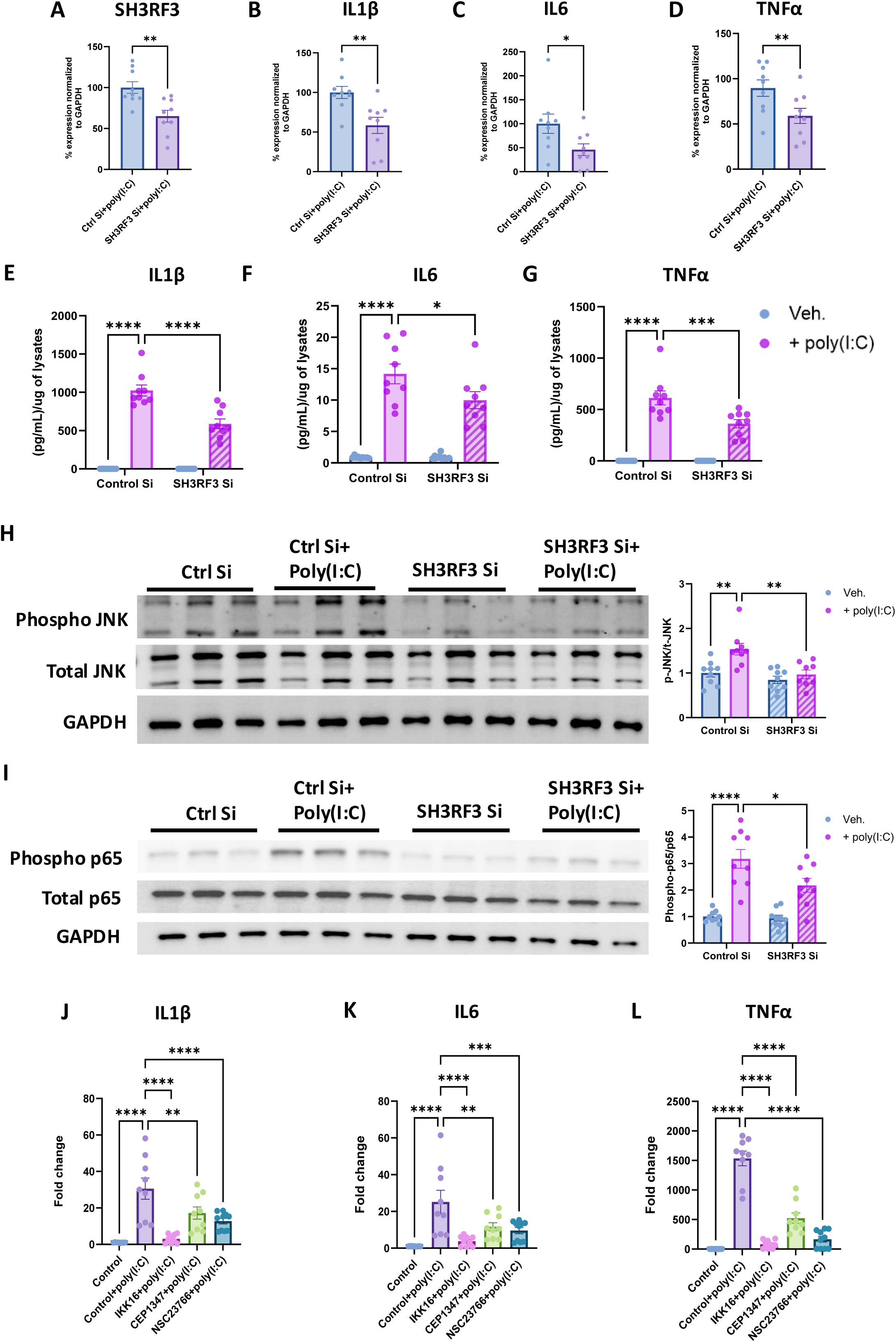
*SH3RF3* knockdown reduces inflammatory cytokine production in iMGLs treated with poly(I:C) via modulation of JNK and NFκB pathways. **(A-D)** iMGLs treated with either *SH3RF3* or control siRNA were subjected to poly(I:C) or vehicle for 2h. qPCR analysis of *SH3RF3* (A), IL1β (B), IL6 (C) and TNFα (D) showed successful knockdown of *SH3RF3* and induction of all cytokines upon poly(I:C) treatment which were significantly reduced by *SH3RF3* KD. Percentage expression was calculated by normalizing gene of interest with respective loading control (GAPDH) for each sample and then all data were normalized to the poly(I:C) treated control siRNA condition and is presented as mean±SEM for N=3 biological replicates per experiment for three independent experiments (N=9). Comparisons between both control Si with poly(I:C) and *SH3RF3* Si with poly(I:C) were made by unpaired t-test, *<0.5, **<0.01. (E-G) Multiplex ELISA analysis of inflammatory cytokines from conditioned media from iMGLs treated with poly(I:C) for 16h. IL1β (E), IL6 (F) and TNFα (G) all show significant induction by poly(I:C) which is reduced by *SH3RF3* KD. Data is presented as mean±SEM for N=3 biological replicates per experiment for three independent experiments (N=9). Statistical comparison among multiple groups were made using two-way ANOVA. (H-I) Western blot analysis of JNK and NFκB pathway activation from iMGLs treated with *SH3RF3* or control siRNA subjected to poly(I:C) for 6 hr. Representative blots for phosphorylated JNK and total JNK (H), phosphorylated p65 and total p65 (I) with densitometry quantification for 3 independent experiments. GAPDH is used as loading control. Data is presented as mean±SEM for N=3 biological replicates per experiment for three independent experiments (N=9). Statistical comparison among multiple groups were made using two-way ANOVA. (J-L) qPCR analysis of IL1β (J), IL6 (K) and TNFα (L) in iMGLs treated with poly(I:C) for 2 hr, with the presence or absence of pathway inhibitors, all of which reduce transcription of all 3 cytokines. CEP-1347 is a JNK pathway (MLK/MAP3K) inhibitor, IKK16 is a NFκB pathway inhibitor, and NSC23766 is a Rac1 inhibitor. Data is presented as mean±SEM for N=3 biological replicates per experiment for three independent experiments (N=9). Statistical comparison among multiple groups were made using one-way ANOVA. *<0.5, **<0.01, ***<0.001, ****<0.0001

We then investigated whether modulation of JNK and NFκB signaling pathways were responsible for the *SH3RF3* KD reduced inflammatory cytokine phenotype in iMGLs. *SH3RF3* KD was performed in presence and absence of poly(I:C) treatment for 6 hours and lysates were subjected to western blot. We observed increased JNK phosphorylation (Thr183 and Tyr185), and hence increased JNK activity, for multiple JNK isoforms upon poly(I:C) treatment (**Figure 3H).** *SH3RF3* KD showed reduced JNK phosphorylation compared to control siRNA poly(I:C) treated conditions. We also assessed if poly(I:C) treatment of iMGLs could induce activation of the p65 (Rel-A) subunit of NF-kB signaling and the effect of *SH3RF3* KD on NF-kB activation. We observed significantly upregulated p65 phosphorylation, demonstrating strong activation of NF-kB signaling, which also was reduced by *SH3RF3* KD. (**Figure 3I**). Similar results were seen for the p105 subunit NFKB1 (**Figure S3F**). In summary, *SH3RF3* levels regulate activity of JNK and NF-kB signaling consistent with these pathways mediating inflammatory response in iMGLs upon poly(I:C) stimulation.

### Pharmacological inhibitors of JNK and NF-kB phenocopy *SH3RF3* KD

To further solidify the mechanistic link between reduced JNK and NF-kB signaling and reduced cytokine production in response to poly(I:C) treatment, we sought to determine if effects of *SH3RF3* KD on inflammatory cytokine production could be mimicked by pharmacological inhibition of these pathways. To inhibit JNK signaling, we employed CEP-1347, a semisynthetic mixed lineage kinase (MLK) inhibitor, which blocks JNK activation by inhibiting upstream MLK family member MAP3Ks^58^. CEP-1347 was used in a clinical trial for Parkinson’s disease, where it failed due to efficacy rather than safety concerns^59^. We also used IKK16, a selective inhibitor of IkB kinase resulting in inhibition of canonical NF-kB signaling^60^. We treated iMGLs with poly(I:C) in presence of CEP-1347, IKK16, or DMSO vehicle control and assessed inflammatory cytokine transcription. CEP-1347 and IKK16 produced significant reduction in transcription of all three inflammatory cytokines, although IKK16 was more potent (**Figure 2J-L**). *SH3RF3* interacts with activated RAC1 upstream to JNK and NF-kB kinases^61^, which led us to hypothesize that inhibition of RAC1 activity might act similarly as *SH3RF3* KD^9^. Treatment with the RAC inhibitor NSC23766 also produced a similar phenotype as *SH3RF3* KD with poly(I:C) stimulated iMGLs, although used at a high concentration (50 μM) due to its low potency. (**Figure 3J-L**)^62^. Taken together, this data demonstrates that inhibition of JNK or NF-kB signaling is sufficient to lower inflammatory cytokine production in iMGLs.

### *SH3RF3* KD reduces inflammatory cytokines produced by iMGLs treated with oligomeric Aβ42 by modulating JNK and NF-kB signaling

We next wanted to extend our findings regarding effects of *SH3RF3* KD on cytokine production to another AD-relevant inflammatory stimuli. Oligomeric amyloid-β (oAβ) peptides are key drivers of AD according to the amyloid hypothesis^1^ and cause microglial activation and release of inflammatory cytokines^63,64^. We utilized a commonly used Aβ oligomerization protocol^65^ which we then assessed for oligomerization state. Coomassie blue staining for total protein showed one dominant band consistent with Aβ trimers and no high order species (**Figure 4A**). Interestingly, western blot using the 6E10 Aβ antibody showed the presence of higher order oligomers (**Figure 4B**), which presumably have greater affinity for the antibody by having multiple epitopes even though a minority product. oAβ42 treatment of iMGLs for 24h resulted in induction of inflammatory cytokines as assessed by qPCR for IL1β, IL6 and TNFα transcripts compared to vehicle treated controls (**Figure 4C-E**). Corroborating with our previous results with poly(I:C) stimulated iMGLs, *SH3RF3* KD produced significantly reduced transcripts for all 3 cytokines.

**Figure 4:**
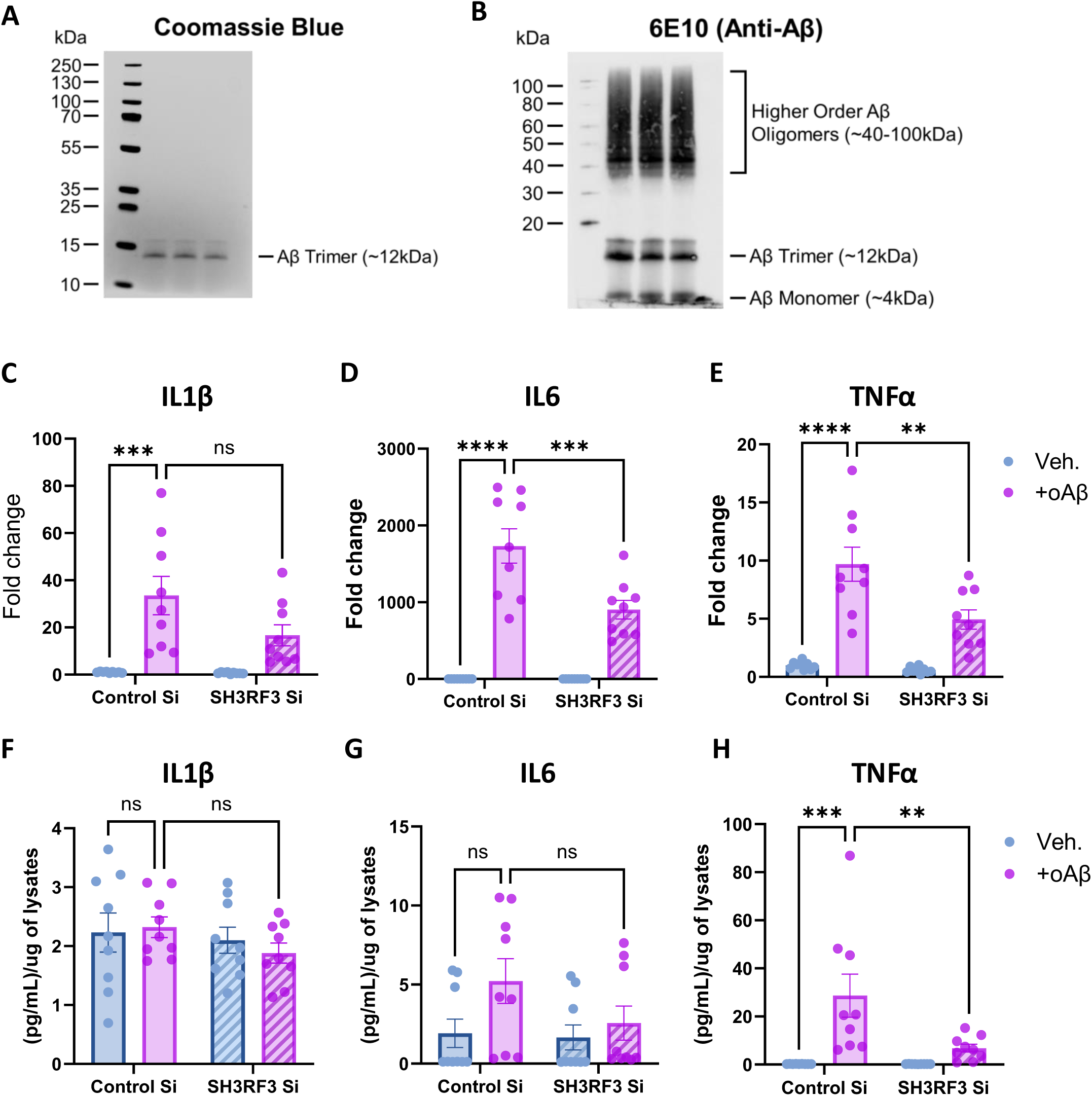
*SH3RF3* knockdown reduces inflammatory cytokines in iMGLs treated with oAβ42. (A-B) Recombinant monomeric Aβ42 was oligomerized overnight and used for experiments the following day. 200ng of oligomers were run on a 4-20% bis-tris gel and stained with Coomassie blue (A) or transferred onto nitrocellulose membrane (B) to probe against amyloid-β (6E10 antibody). The majority of oligomers have a size consistent with trimers, although higher order species are also present. (C-E) qPCR analysis of inflammatory cytokines IL1β (C), IL6 (D) and TNFα (E) in iMGLs treated with *SH3RF3* or control siRNA subjected to 5 μM oAβ42 or vehicle for 24h. Inflammatory cytokines were statistically reduced by *SH3RF3* KD, data is presented as mean±SEM for N=3 biological replicates per experiment for three independent experiments (N=9). Statistical comparison among multiple groups were made using two-way ANOVA. (F-H) Multiplex ELISA analysis of inflammatory cytokines IL1β (F), IL6 (G) and TNFα (H) from conditioned media from iMGLs treated with *SH3RF3* or control siRNA subjected to 5 μM oAβ42 for 3 hr. TNFα is statically reduced by *SH3RF3* KD. Data is presented as mean±SEM for N=3 biological replicates per experiment for three independent experiments (N=9). Statistical comparison among multiple groups were made using two-way ANOVA. **<0.01, ***<0.001, ****<0.0001.

Although we initially started with 24h treatment as a point of analysis, we realized that oAβ42 would also be actively phagocytosed by iMGLs which would reduce the potency of the stimulus over time. We hypothesized this could at least partially explain a recently published study which only showed minor transcriptional effects of iMGLs treated for 24 hours with oAβ42^66^, although that study also used a lower concentration of oAβ42 (1μM instead of 5μM) and potentially had a different Aβ oligomerization state. Thus, we did a time course experiment, which showed transcriptional responses at 24h to oAβ42 were already much lower than at earlier 6h and 12h time points (**Figure S4A**). We also determined that NF-kB signaling was activated by 3h as assessed by western blot and was more robust for 5μM as compared to 1μM oAβ42 (**Figure S4B**).

Upon learning that oAβ42 transcriptional effects were blunted by 24h post-treatment, we tested if iMGLs treated with oAβ42 for 3h released inflammatory cytokines and if *SH3RF3* KD affected this response. While IL1β levels were not increased by oAβ42 at this time point, IL6 and TNFα levels were significantly elevated (**Figure 4F-G**). *SH3RF3* KD significantly reduced TNFα release and trended towards reducing IL-6. We also evaluated effects of oAβ42 on JNK and NF-kB pathways in the presence of *SH3RF3* KD. As anticipated, oAβ42 activated JNK and NF-kB pathways as observed by increased phosphorylation of JNK and p65 (**Figure 5A-B**) as well as p105 (**Figure S5A**). Similar to poly(I:C) treatment, *SH3RF3* KD reduced activation of JNK and NF-kB signaling. We also performed similar chemical inhibitor experiments which demonstrated that JNK inhibition by CEP-1347 or NF-kB by IKK16, were sufficient to block inflammatory cytokine production by oAβ42 in iMGLs (**Figure 5C-E**). In summary, inflammatory cytokines produced by iMGLs treated with oAβ42 are also attenuated by *SH3RF3* KD, which is likely mediated by reduced activation of JNK and NF-kB signaling.

**Figure 5:**
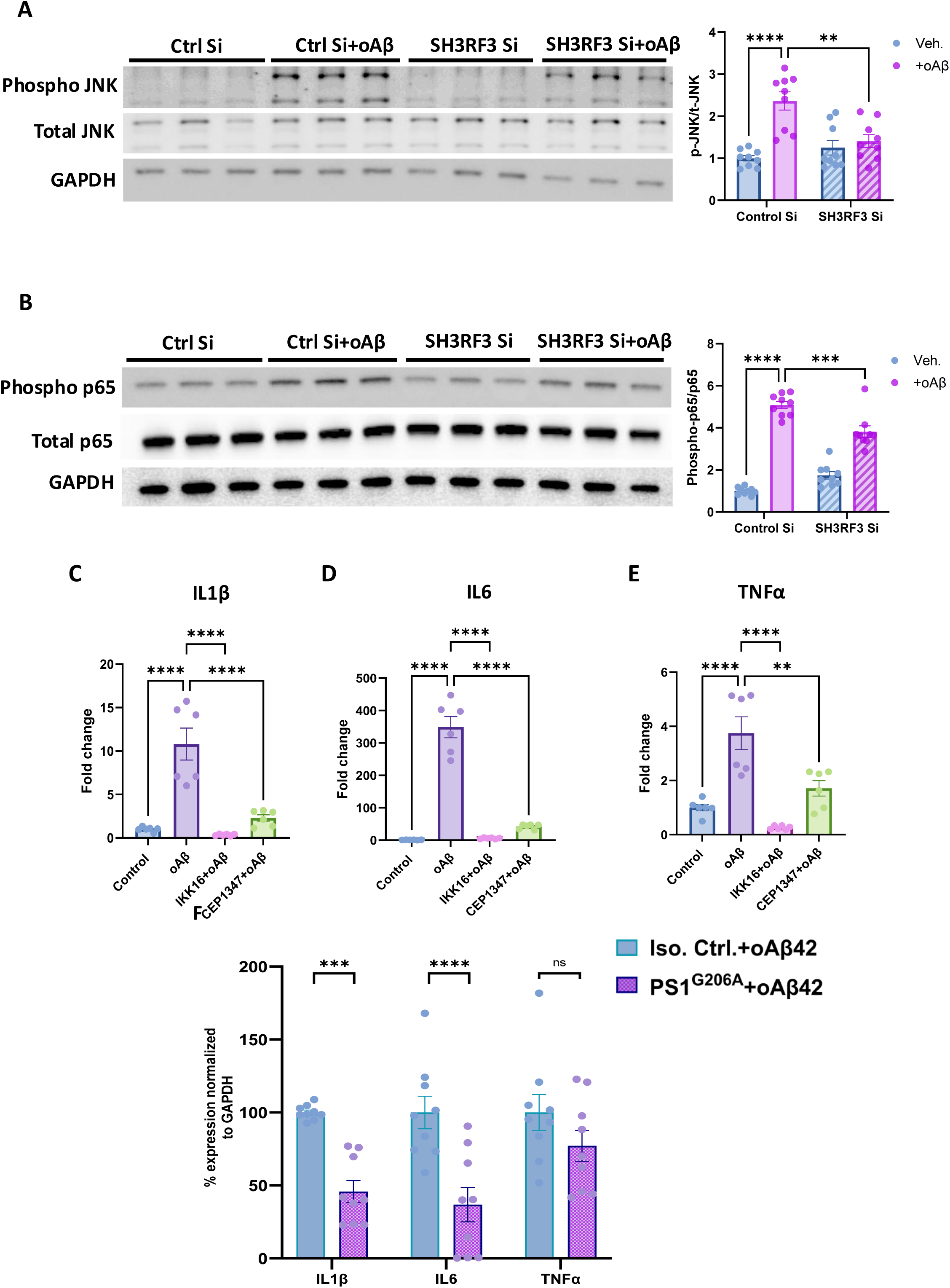
*SH3RF3* KD reduces JNK and NFkB pathway activation induced by oAβ42 treatment in iMGLs, which is sufficient to reduce inflammatory cytokine production. (A-B) Representative immunoblots and densitometry analysis of JNK (pJNK/JNK, A) and NFkB (phosho-p65/p65, B) pathway activation in iMGLs treated with *SH3RF3* or control siRNA subjected to oAβ42 or vehicle for 3h. Densitometry data is presented as mean±SEM for N=3 biological per experiment for three independent experiments (N=9). Statistical comparison among multiple groups were made using two-way ANOVA. (C-E) qPCR analysis of inflammatory cytokines IL1β (C), IL6 (D) and TNFα (E) upon 5 μM oAβ42 treatment for 12h in iMGLs pre-treated with either DMSO vehicle, CEP-1347 (JNKi), or IKK16 (NFkBi). Data is presented as mean±SEM for N=3 per experiment for two independent experiments (N=6). Statistical comparison among multiple groups was made using one-way ANOVA. (F-H) qPCR analysis of inflammatory cytokines IL1β (F), IL6 (G) and TNFα (H) upon 5 μM oAβ42 treatment or vehicle for 6h in isogenic control and *PSEN1*^G206A^ iMGLs. Data is presented as mean±SEM for N=3 biological replicates per experiment for three independent experiments (N=9). Statistical comparison among multiple groups were made using two-way ANOVA. **<0.01, ***<0.001, ****<0.0001.

### *PSEN1*^G206A^ microglia have reduced induction of cytokines in response to oligomeric Aβ42

We hypothesized that *SH3RF3* protective SNPs might be particularly beneficial in *PSEN1* mutation carriers due to their strong effect on age on onset in that *PSEN1*^G206A^ (**Table 2**). It has been previously reported that *PSEN1*^ΔE^^9^ iMGLs have reduced inflammatory responses to bacterial lipopolysaccharide (LPS) relative to isogenic control iMGLs.^67^ To test if *PSEN*^G206A^ iMGLs also have reduced inflammatory responses, we knocked the G206A mutation into both alleles of *PSEN1* in the IMR90 (cl.4) hiPSC line using CRISPR/Cas9 (**Figure S5B**). Homozygous rather than hemizygous knock-in was confirmed via qgPCR (data not shown)^68^. *PSEN1*^G206A^ and isogenic control iMGLs were then treated with oAβ42 for 6h to look for transcriptional effects on IL1β, IL6 and TNFα (**Figure 5F**). While the level of transcriptional response to oAβ42 were more variable between independent experiments using these lines, IL1β and IL6 were consistently reduced in *PSEN1*^G206A^ iMGLs compared to isogenic control iMGLs. TNFα trended lower. This suggests that microglia in subjects with *PSEN1* mutations may have reduced pro-inflammatory responses in general.

## Discussion

It is critical to understand the mechanisms of why some high-risk individuals are protected from getting AD as an alternative therapeutic strategy to targeting neuropathological changes in amyloid and tau directly. In this study, we focused on *SH3RF3*, which we previously identified as protective against AD in *PSEN1*^G206A^ carriers^4^. Our discovery dataset of *PSEN1*^G206A^ carrier families yielded a set of contiguous SNPs with a varying degree of delay in age at onset. Then the subsequent examination of familial and sporadic LOAD in Caribbean Hispanics (using *APOE4* as a risk variant) confirmed the modifying effects of *SH3RF3,* in which the significant association was observed in the presence of *APOE4*, but not in the absence of *APOE4*. We further note that these variants were located in gene enhancer regions, supporting observed varying degree of gene expressions.

Thus, we then sought to understand the mechanisms of how a protective SNP in *SH3RF3*^4^ delayed onset of AD in *PSEN*^G206A^ carriers. We focused on JNK and NF-kB signaling since SH3RF family members including *SH3RF3* had been shown to be able to activate these pathways^9–12^. For JNK signaling, SH3RF proteins (as best described for SH3RF1/POSH) act as scaffolds linking upper and lower components of this MAPK cascade^12,13^. The mechanism(s) of how *SH3RF3* modulates NF-kB activity is not well understood and should be the subject of further research. We initially focused on neurons, as early-onset AD mutations like those in *PSEN1* are primarily thought to drive disease through modulating APP processing, such as by increased Aβ42:Aβ40 ratio. However, *SH3RF3* KD did not affect APP processing in a significant fashion under steady-state conditions (**Figure 2**). *SH3RF3* KD did decrease phosphorylation at the JNK-specific tau S422 phospho-site which could have some effect on AD risk. It is important to note that while phosphorylation of the S422 site of tau is found in PHFs in AD brains^37^ and has been proposed to be pathogenic^69^, it also been reported that phosphorylation of tau at S422 blocks caspase 3 cleavage of tau^70^, which is considered a pro-pathogenic event. In addition, one significant caveat of our neuronal studies is that as JNK and NF-kB are stress-induced signaling cascades, thus, *SH3RF3* may play a stronger role in neurons in response to specific stresses not tested here.

We then chose to look at microglia as JNK and NF-kB signaling are also critical for inflammatory responses in this cell type^43,44^. We chose to focus our analysis on 3 pro-inflammatory cytokines, IL1β, IL6 and TNFα, which have been associated with AD and are members of a human-specific microglial cytokine response cluster (CRM2) activated in human microglia xenotransplanted into the humanized APP^NL-G-F^ AD mouse model^30^. *SH3RF3* KD can blunt pro-inflammatory cytokine production from two AD-relevant stresses, a viral mimic poly(I:C) and oAβ42 (**Figures 3-5**). This was associated with reduced JNK and NF-kB signaling, and blockade of these pathways through chemical inhibitors was sufficient to reduce inflammatory cytokine production. Overall, our data is consistent with a model where reduced expression of *SH3RF3* by protective SNPs lowers inflammatory cytokines produced by inflammatory insults by dampening JNK and NF-kB signaling, to consequently delay the onset of AD (see **Graphical Abstract**). Interestingly, while genes expressed primarily in microglia have almost exclusively been associated with late-onset forms of AD, and early-onset AD has been associated with neuronal acting genes, our study suggests that early-onset AD risk may also be significantly influenced by genetic modulators of microglial function. Why microglia from *PSEN1* mutation carriers (ΔE9^67^ or G206A (**Figure 5**)) have reduced inflammatory responses remains unclear. As TREM2 is a direct γ-secretase target^71^, an intriguing possibility is that *PSEN1* mutant microglia have reduced/altered γ-secretase activity against TREM2 which leads to enhanced TREM2 intracellular signaling and reduced inflammatory processes, which could be investigated in future studies.

One limitation of our study is that we focused exclusively in microglia on inflammatory cytokine production. Other inflammatory responses such as ROS production or effects on phagocytosis should be addressed in future studies. Regarding inflammatory reaction of iMGLs oAβ42, it is important to note that this is a very transient response that could reflect elevated oAβ from transient pathological events like ischemia which increase Aβ levels^72^, rather than a more “DAM”-like response to aggregated fibrillar forms of Aβ^73^. On the other hand, xenotransplantation of iMGLs into the APP^NL-G-F^ AD mouse model^30^ had human-specific cytokine modules under both steady-state responses to amyloid and co-injection of oAβ42, suggesting a broader role of inflammatory cytokine production in the human microglial response to AD pathology. Importantly, *SH3RF3* was a member of the CRM2 module suggesting it could help drive this cluster which could be tested in future xenotransplantation experiments.

Finally, a key goal of trying to understand the molecular mechanisms of AD protective SNPs was to mimic those genetic protective mechanisms with small molecules. We found CEP-1347, a MAP3K inhibitor that blocks JNK signaling^58^, significantly reduced inflammatory cytokine production in iMGLs from both a viral mimic and oAβ42. CEP-1347 and other MAP3K inhibitors have previously shown benefits in blunting inflammatory responses in other microglial/macrophage models and promote Aβ clearance^26,74,75^. Importantly, CEP-1347 has also been shown to block neuronal cell death from oAβ42, trophic factor withdrawal, UV, and oxidative stress^25,76^. This data led, at least in part, to CEP-1347 being tested in the PRECEPT trial for Parkinson’s disease (PD), where it failed for efficacy rather than safety concerns^77^. One possible reason for this failure was that neuronal cell death in PD has been reported to be driven by the MAP3K DLK^78^, which CEP-1347 inhibits to a much lesser degree than MLKs^58^. In addition, genetic evidence suggests that microglia play a more dominant role in AD than PD, and therefore, CEP-1347 effects on microglial inflammatory responses would be expected to have greater importance in AD. Thus, reconsideration of using MAP3K (MLK) inhibitors like CEP-1347 to treat AD may be warranted as this could act as a double hit to both dampen pro-inflammatory microglial responses and to protect neurons from cell death from Aβ and other insults. This particularly could be the case for *PSEN1* mutant carriers, as we (**Figure 4F**) and others^67^ have seen a lower baseline inflammatory response in *PSEN1* mutant microglia, as well as *PSEN1* mutant carriers having a more clearly Aβ-driven form of AD.

## Acknowledgements

This work was primarily supported by NIA R01AG058918 (J.L.H PI, AAS Co-I) and R56 AG051876 (J.H.L). It was also supported by BrightFocus (A2015633S; J.H.L.), 1P30AG066462-01 (A.A.S. Co-PI on Development grant), the Henry and Marilyn Taub Foundation (A.A.S.), and the Thompson Family Foundation (TAME-AD, A.A.S.). Data used in the replication analyses were supported by R01AG067501 (R.M.) and U24AG056270 (R.M.)

## Author contributions

R.P., R.C., C.L.C., G.S., E.T., S.M. and A.A. conducted experiments and analyzed data. A.P., A.M. and I.J-V., contributed to the collection and analysis of *PSEN1* and control subject samples. R.M. provided additional data sets for replication of the findings from the initial analysis. R.P., R.C. J.H.L, and A.A.S. designed experiments and wrote the manuscript. All authors edited the manuscript. A.A.S. and J.H. L. supervised all research.

## STAR Methods

### CHART OF ALL REAGENTS

#### Chart of All Reagents

| Reagent or Resource | Source | Identifier |
| --- | --- | --- |
| **Antibodies** |  |  |
| p-SAPK/JNK (T138/Y185) (81E11) | Cell Signalling Technologies | cat.# 4668S; RRID: AB_823588 |
| SAPK-JNK | Cell Signalling Technologies | cat.# 9252S; RRID: AB_2250373 |
| p-NF-kappaβ P105 (S932)(18E6) | Cell Signalling Technologies | cat.# 4806S; RRID: AB_2282911 |
| NF-kappaβ P105/P50 (SD10D11) | Cell Signalling Technologies | cat.# 13681S |
| p-c-JUN (S73) (D47G9) | Cell Signalling Technologies | cat.# 3270S; RRID: AB_2895041 |
| c-JUN (L70B11) | Cell Signalling Technologies | cat.# 2315S |
| CD45 Alexa fluor 647 conjugated | Biolegend | cat.# 304056; RRID: AB_2564155 |
| CD11b Alexa fluor 488 conjugated | Biolegend | cat.# 301318; RRID: AB_493017 |
| IBA1 | FUJIFILM | cat. # 019-19741 |
| P2RY12 | Millipore-Sigma | cat.# HPA014518; RRID: AB_2669027 |
| TREM2 | Cell Signalling Technologies | cat.# 91068S; RRID: AB_2721119 |
| Anti-Aβ 6E10 | BioLegend | cat.# SIG-39320; RRID: AB_2715854 |
| Phospho-APP (Thr668) | Cell Signalling Technologies | cat.#6986; RRID:AB_10831197 |
| APP (Y188) | Abcam | cat.# ab_32136; RRID_2289606 |
| Phospho-Tau (Thr217) | Abcam | cat.#ab_192665;RRID_2139718 |
| Phospho-Tau (ser422) | Thermofisher | Cat.#44-764G; RRID_AB_2533748 |
| Total Tau | Dako | Cat.# A0024;RRID_AB_10013724 |
| GAPDH | Thermofisher | cat.# MA1-16757; RRID: AB_568547 |
| **Chemicals, peptides, and recombinant proteins** |  |  |
| IL34 | R&D | cat.# 5265-IL-010/CF |
| TGFβ | R&D | cat.# 240-B-010/CF |
| m-CSF | R&D | cat.# 216-MC-010/CF |
| CD200 | Bon-Opus Bio | cat.# BP004 |
| CX3CL1 | R&D | cat.# 365-FR/CF |
| CEP1347 | Tocris | cat.# 4924 |
| IKK16 | Tocris | cat.# 2539 |
| NSC23766 (RAC1 inhibitor) | EMD Millipore | cat.# 553502 |
| Amyloid-β-1-42 | rPeptide | cat.# A-1163-2 |
| Poly(I:C) | Tocris | cat.# 4287 |
| STEMdiff Hematopoietic Kit | STEMCELL Technologies | cat.# 05310 |
| DMEM-F12 medium | Thermo Fisher | cat.# 11320033 |
| Non-essential amino acids | Thermo Fisher | cat.# 11140050 |
| Insulin-transferrin-selenium | Thermo Fisher | cat.# 41400045 |
| Insulin | Sigma-Aldrich | cat.# 12585014 |
| Glutamax | Thermo Fisher | cat.# 35050061 |
| B27- supplements | Thermo Fisher | cat.# 17504044 |
| N2- supplements | Thermo Fisher | cat.# 17502048 |
| 1-Thioglycerol | Sigma-Aldrich | cat.# M-6145 |
| Pen Strep | Thermo Fisher | cat.# 15140122 |
| SuperSignal West Pico PLUS | Thermo Fisher | cat.# 34577 |
| **Critical commercial assays and kits** |  |  |
| Aβ peptide panel 1 ELISA kit | Mesoscale Discovery | cat. # K15200E |
| U-Plex biomarker group 1<br>(human) ELISA kit | Mesoscale Discovery | cat. # K15067L |
| Genomic cleavage assay kit | GeneArt, ThermoFisher | cat. #A24372 |
| RNeasy Plus Micro Kit | Qiagen | cat.# 74034 |
| BCA Protein Assay Kit | Millipore Sigma | cat.# 71285-3 |

| Experimental models: Cell lines |  |  |
| --- | --- | --- |
| H9 hESC | WiCell | WA09 NIH registry |
| IMR90 cl.4 iPSC | WiCell |  |
| FA0000010 (FA10) hiPSC | RUCDR |  |
| HEK293 | ATCC |  |
| Plasmids |  |  |
| pMLM3636 | Addgene | cat.# 43860, RRID:Addgene_43860 |
| eSpCas9(1.1) | Addgene | cat.# 71814, RRID:Addgene_71814) |

#### Primers

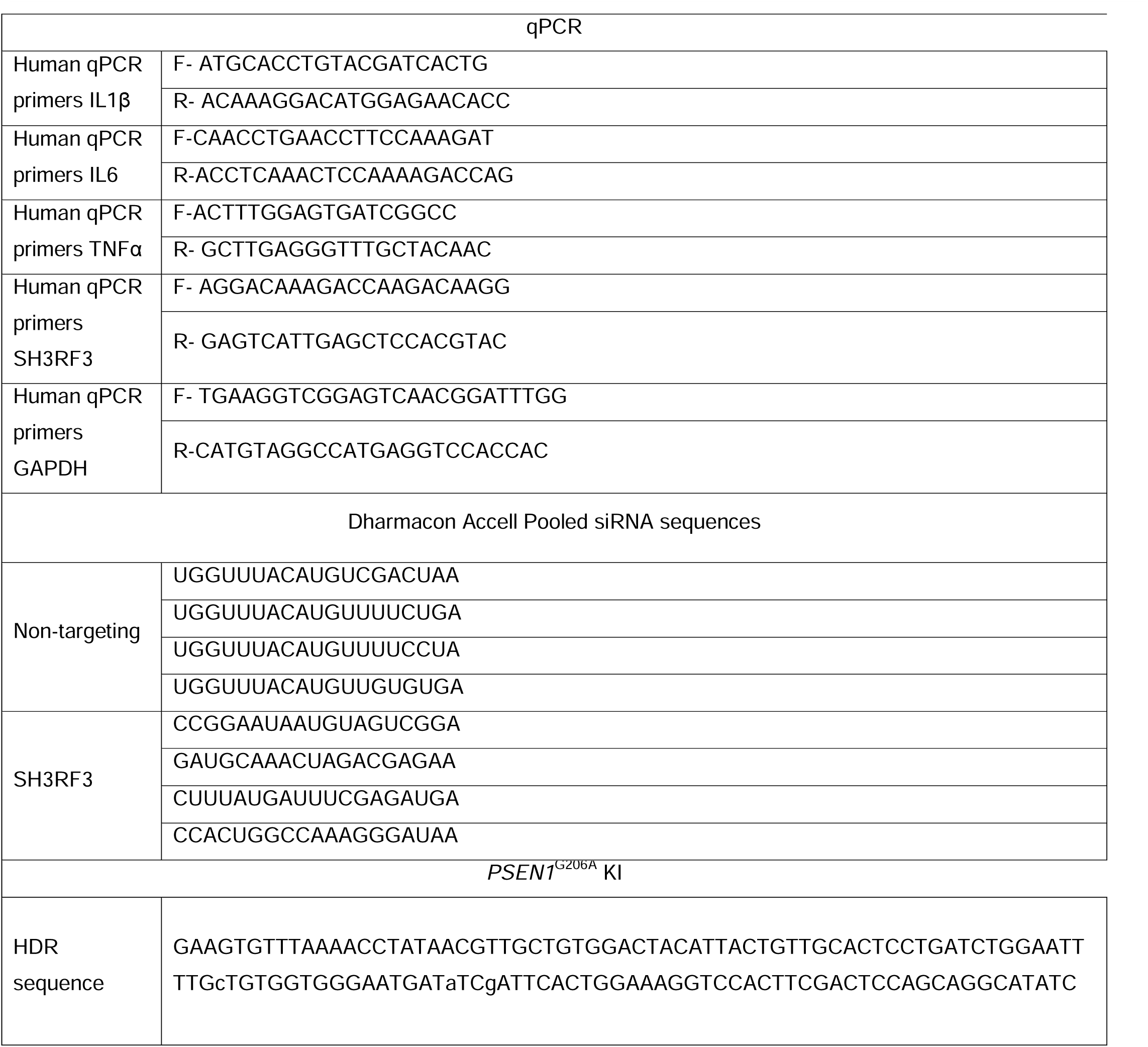
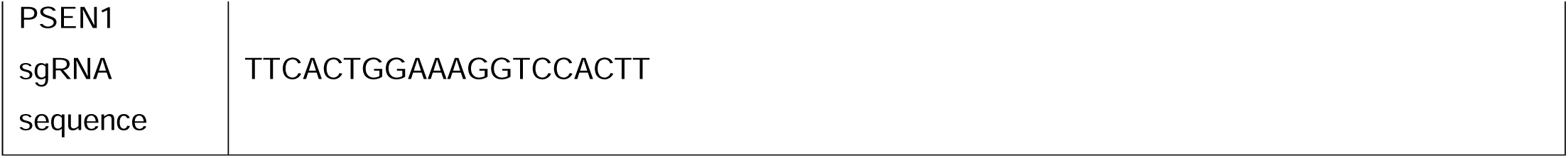

##### Genetic association analysis Extending

our earlier work^4^, we performed WGS, rather than whole exome sequencing, on a larger set of Caribbean Hispanic families with at least one PSEN1^-G206A^ carrier. The study protocol was approved by the Institutional Review Board of Columbia University and the University of Puerto Rico, School of Medicine, and written informed consent was provided by all study participants. To confirm the findings from our discovery dataset and generalize to more common forms of AD (i.e., familial and sporadic LOAD), we examined 1,946 singletons and 2,114 family members of the familial late onset AD (LOAD) comprised of 4,060 Caribbean Hispanic elderly, primarily from the Dominican Republic (EFIGA^31^). In addition, we examined 2,058 Hispanic elderly with primarily sporadic AD (WHICAP^32^).

###### Whole genome sequencing (WGS)

To enhance the resolution of the *SH3RF3* gene, we first performed WGS on about two-third of the family members (n=337) from a total of 505 family members in 72 families. The genome sequencing used one μg of DNA, an Illumina PCR-free library protocol, and sequenced on the Illumina HiSeq platform at the NY Genome Center. We assessed sex mismatches and unexpected familial relationships prior to analyses. Genomes were sequenced to a mean coverage of 30X. Sequence data analysis was performed using an automated analysis pipeline that matched the Centers for Common Disease Genomics (CCDG) and TOPMed recommended best practices. Briefly, sequencing reads were aligned to the human reference, hs38DH, using Burrows-Wheeler Alignment - Maximal Exact Matches (BWA-MEM) version 0.7.15. Variant calling was performed using the Genome Analysis Toolkit (GATK) best practices. Variant filtration was performed using Variant Quality Score Recalibration (VQSR) at tranche 99.6%, which identifies annotation profiles of variants that could be called with high confidence and assigns a score Variant Quality Score Log-Odds (VQSLOD) to each variant. Variants passing the VQSR threshold were further filtered out for sample missingness (>2%), depth of coverage (DP) <10 and genotype quality (GQ) >20. We then annotated high-quality variants using Annotate Variation (ANNOVAR). Specifically, variants were annotated for population-level frequency using Genome Aggregation Database (gnomAD), in silico function using Variant Effect Predictor (VEP), and variant conservation using Combined Annotation Dependent Depletion (CADD) score.

###### Genome-Wide Association (GWA) Analysis

For approximately one-third of the family members, we performed GWA using the Illumina Human Hap 650k and Illumina 1M arrays and Illumina Infinium General Screening Array V2 or V3 (Illumina, San Diego, CA, USA) with added disease markers at the Center for Applied Genomics at Children’s Hospital of Pennsylvania. Genotype calling was done using Illumina’s GenomeStudio v2.0.5 with the Genotyping module; in total, 664,337 QCed variants were genotyped for 430 study participants from *PSEN1*^G206A^ Cohort, and 572,421 QC’ed variants for 4,060 samples of EFIGA cohort and 532,147 QCed variants for 2,058 samples of WHICAP cohort.

###### Imputation

To increase the coverage of the genome, we performed imputation on all autosomal chromosomes using the TOPMed Imputation Server (https://imputation.biodatacatalyst.nhlbi.nih.gov/), and the TOPMed Reference Panel (Version R2) built on whole genome sequencing data from 97,256 individuals and 308,107,085 genetic variants of diverse ancestral and ethnic backgrounds. This TOPMed panel included 487 Puerto Ricans, 1,062 Costa Ricans and 504 Mexicans to enhance the relevance of the reference panel (TOPMED ref). A previous study had shown that imputation in these Puerto Ricans was accurate^79^. We included all variants with imputation quality of at least R^2^ >0.30 for further analysis.

Prior to analysis, we conducted the standard quality control checks, including checking of the reported family relationships using the software PREST (https://www.utstat.utoronto.ca/sun/Software/Prest/), Hardy-Weinberg equilibrium, and genotyping rate. The details of the quality control procedures are presented elsewhere^80^.

###### Statistical Analysis

We first examined the relationship between genetic variants and age at last follow up (i.e., age at onset for affected family members and age at last examination for unaffected family members) using 337 family members with WGS data. This analysis applied a mixed linear model, adjusting for sex, *APOE4* genotype, *PSEN1*^G206A^ status, affection status, principal components (PC1-PC3), and genetic relationship matrix (GRM). PC1 to PC3 were included to adjust for genetic background, and GRM was included to take into account relatedness among family members. Subsequently, we re-analyzed the data after adding 168 family members who lacked WGS data but had imputed genotype data. Our earlier work showed that imputation in Caribbean Hispanic families was accurate^79^. We applied this 2-stage approach to gain finer resolution, while maintaining cost-effectiveness. We note that the same set of models were performed in EFIGA and WHICAP replication datasets; however, WHICAP did not require an inclusion of GRM since it was not a family-based study.

##### hPSC maintenance and differentiation into neurons and microglia-like cells (iMGLs)

The FA0000010 (FA10) hiPSC line was acquired from RUCDR, and the IMR90 (cl.4) hiPSC line^81^ and the H9 (WA09) hESC line from WiCell. All cultures were grown in the presence of penicillin/streptomycin unless noted otherwise (ThermoFisher) Undifferentiated hPSCs were maintained in StemFlex media (ThermoFisher,) on reduced growth factor Cultrex BME (Biotechne), and routinely split 1-2 times a week with ReLeSR (STEMCELL Technologies) without ROCKi (Y-27632, Selleck Chemicals) as described previously.^33^ Generation and transdifferentiation of permanent transgene hPSCs(Ngn2/rtTA) into excitatory neurons was described in detail previously.^33^ In brief, undifferentiated H9 hPSCs with permanent Ngn2/rtTA transgenes (via puromycin selection) were plated in presence of 10 mM Y-27632 (ROCKi) on PEI/Laminin-coated surfaces at 55,000-60,000 cells/cm^2^ (200k/well of a 12 well) in N2/B27 medium supplemented with 1 mg/mL doxycycline (Sigma-Aldrich), 10 ng/mL BDNF (R&D systems), 10 ng/mL NT3 (R&D systems), and 2 uM AraC (cytarabine, Tocris) which was added for first 7 days of differentiation as needed to deplete non-neuronal cells from culture. At day 7, cultures were switched to Brainphys (N2/SM1) neuronal differentiation medium (STEMCELL Technologies) supplemented with 1 mg/mL doxycycline, 10 ng/mL BDNF, 10 ng/mL NT3 with half medium change every other day for the duration of differentiation. Brainphys neuronal differentiation medium was further supplemented with 10% mouse astrocyte conditioned media (ACM; ScienCell) from d14 until samples are collected on d28.

hiPSCs were differentiated into iMGLs as described previously with minor adaptations.^57,82^ hiPSCs were differentiated into hematopoietic precursors cells (HPCs) using the STEMdiff Hematopoietic kit (STEMCELL Technologies) largely as per manufacturer’s instructions. In brief, on day −1 iPSCs were detached with ReLeSR and passaged to achieve a density of 1–2 aggregates/cm^2^ of 100-150 cells. Multiple densities were plated in parallel. On day 0, colonies of appropriate density were switched to Medium A from the STEMdiff Hematopoietic Kit to initiate HPC differentiation. On day 3, cells were switched to Medium B with a full media change and fed again with a full media changed on day 5. Cells remained in Medium B for the rest of the HPC differentiation period with Medium B overlay feeds every other day. HPCs were collected 3 independent times by gently removing the floating population with a serological pipette at days 11, 13 and 15 (or days 12, 14 and 16). HPCs were either cryopreserved in 45% Medium B, 45% knockout serum replacement (ThermoFisher) and 10% DMSO and stored in liquid nitrogen or directly plated for iMGL induction. Pilot differentiations showed a majority of HPCs were CD43^+^ as measured by flow cytometry (>90%). HPCs were terminally differentiated by plating 35,000-60,000 cells/cm^2^ HPCs on reduced growth factor Cultrex BME in microglia medium (DMEM/F12, 2X insulin-transferrin-selenite, 2X B27, 0.5X N2, 1X glutamax, 1X non-essential amino acids, 400 mM monothioglycerol, and 5 mg/mL human insulin (ThermoFisher) freshly supplemented with 100 ng/mL IL-34, 25 ng/mL M-CSF (R&D Systems) and 50 ng/mL TGFβ1 (STEMCELL Technologies) for every other day until day 24. For the majority of differentiations, differentiating cells were “split” by taking remaining non-adherent cells and plating on to a fresh Cultrex-coated well of the same format (1:1). On day 25, 100 ng/mL CD200 (Bon Opus Biosciences) and 100 ng/mL CX3CL1 (R&D Systems) were added to microglia medium to mimic a brain-like environment. For all experiments in this study, iMGLs were treated between day 28 to 35 (post HPC) and samples were collected on days 34-36. The majority of iMGL experiments were generated from FA10 hiPSCs unless stated otherwise.

##### *SH3RF3* knockdown in neurons and iMGLs

*SH3RF3* knockdown was performed using Accell SMARTpool siRNAs targeting *SH3RF3*, with Accell human non-targeting pool siRNA used as a negative control (Dharmacon/Horizon Discovery). siRNAs were resuspended and delivered as per manufacturer’s instructions. In brief, 50 nmol siRNA was resuspended using 1X siRNA buffer at 100 uM stock concentration and directly delivered at 1 uM concentration in cell culture media (at day 21 for neurons, day 28 for iMGLs). Knockdown was performed for 7-days with media overlay ay day 3 and 5. RNA, protein lysates, and conditioned media were collected from neurons at day 28 and iMGLs at day 35.

##### Treatment of neurons and iMGLs with chemical inhibitors and inflammatory stimuli

On day 28 neurons were treated with okadaic acid or vehicle for 3 hours before lysing for protein and conditioned media. In some experiments, iMGLs were pre-treated with chemical inhibitors or DMSO vehicle for 2 hours before inflammatory stimuli, which included CEP-1347 (100 nM; Tocris), IKK16 (1 uM; Tocris), and NSC23766 (50 uM; EMD Millipore). iMGLs were then treated with either the dsRNA viral mimic poly(I:C) at (25 ug/mL)^83^, oligomerized Aβ42 (5 uM; rPeptide), or vehicle control (dH_2_O or F12 media respectively). Recombinant Aβ42 was oligomerized as described previously, ^84^ and validated via Coomassie blue staining and western blot using the 6E10 antibody (1:1000; BioLegend).

##### qPCR

RNA was extracted using the RNeasy Plus Micro Kit (Qiagen) and resuspended in 30 µl of RNAse-free water. RNA quantity and quality were assessed with a Nanodrop One (ThermoFisher). For qPCR, 100 ng of RNA was transcribed into complementary DNA Maxima First Strand cDNA Synthesis Kit (ThermoFisher). Gene expression was assessed using FAST SYBR^TM^ Green Master Mix (ThermoFisher) using primer sets for target genes on a QuantStudio 7 Flex system (Applied Biosystems) and Bio-Rad CFX Opus 384. Gene expression for target genes are normalized to GAPDH and fold change are calculated as ΔΔC_t_.

##### Amyloid-β42 monomerization, oligomerization and characterization

Amyloid-β42 was monomerized, stored and later oligomerized as previously described.^84,85^ In brief, Aβ(1-42) (rPeptide, 1 mg) vial was allowed to equilibrate to room temperature. Peptides were resuspended in ice-cold Hexafluoroisopropanol (Sigma-Aldrich) to achieve 1 mM concentration directly in a vacuum-sealed vial using Hamilton syringe. The suspension was divided into 8 equal aliquots by volume using a Hamilton syringe into low protein binding Eppendorf tubes, incubated at room temperature for 2h and dried on speedvac at 800g. Upon drying monomerized Aβ(1-42) films can be stored at −80° C for months to years in sealed tubes. For generating oligomers, Eppendorf tubes containing Aβ(1-42) peptide films were equilibrated to room temperature, resuspended in DMSO to achieve 5 mM concentration and sonicated at 17° C for 10 min for complete resuspension. Oligomerization was induced by adding phenol-red free F12 to achieve final concentration of 100 μM and incubated at 4° C for 24 to 48 hours. Recombinant monomeric amyloid-β-42 was oligomerized overnight and used for experiments the following day. 200ng of oligomers were prepared in the absence of reducing reagents and run on a 4-20% bis-tris gel in MES running buffer. Gels were stained with Coomassie blue overnight or transferred onto 0.2um nitrocellulose membrane to probe against amyloid-β (6E10 antibody, BioLegend).

##### Western blotting

iMGLs or neurons were harvested using RIPA Lysis Buffer (ThermoFisher) supplemented with Halt protease and phosphatase inhibitors single-use cocktail (ThermoFisher) on ice. 10 ug of lysates and 1x Laemmli sample buffer/5% BME were boiled at 95 °C for 10 min. SDS-PAGE was performed by loading samples on Bolt 4–12% gradient Bis-Tris gels (ThermoFisher) using Bolt MOPS buffer (ThermoFisher). For western blotting, nitrocellulose membranes were first blocked with Superblock TBS buffer/0.1% Tween 20 (ThermoFisher) for 1h, and then incubated overnight at 4 °C with the following primary antibodies diluted in Superblock/0.1% Tween 20: p-JNK (1:1000; Cell Signaling Technologies), total-JNK(1:1000; Cell Signaling Technologies), phospho-p-50/p105 (1:1000; Cell Signaling Technologies), total p50/p105(1:1000; Cell Signaling Technology), phospho-Tau Ser422 (1:1000; Sigma), phospho-Tau Thr217 (1:1000; Abcam) and GAPDH (1:5000; ThermoFisher). After 3 washes with TBS/0.1% Tween 20, membranes were then incubated with the appropriate HRP-conjugated secondary antibody (1:5000; ThermoFisher) for 1h and washed 3 more times with TBS/0.1% Tween 20. Pierce ECL western blotting substrate (ThermoFisher) or super signal west pico plus chemiluminescent substrate (ThermoFisher) was used to develop chemiluminescence and blots were imaged using the Bio-Rad Chemidoc MP system.

##### Multiplex ELISA for Aβ or lL-1β, IL6 and TNFα

Conditioned media (CM) from neurons or iMGLs were first spun at 1000 rpms for 3 min., and SNF was aliquoted and stored at −80°C. CM was diluted 1:2 in diluent 35 and processed for ELISA as per the manufacturer’s instructions. The Aβ peptide panel 1 ELISA kit and custom cytokine kits U-Plex biomarker group 1 (human) ELISA kit were from Mesoscale Discovery (MSD Quickplex SQ120). Analyte concentrations (pg/mL) was calculated as per manufactures instruction.

##### Immunocytochemistry

iMGLs were plated on low growth factor Cultrex-coated 24-well plates, cultured for 35 days and fixed with 4% paraformaldehyde for 20 min on ice. After fixation, cells were blocked with Superblock blocking buffer in PBS/LJ0.1% Triton-X for 1LJh and then incubated with the following primary antibodies overnight at 4 °C: α-CD11b (1:100; BD Biosciences) and α-Iba1 (1:200; Wako). Next day, cells were washed thrice with PBS/0.1% Triton-X and then incubated with corresponding Alexa Fluor-conjugated secondary antibodies (1:500; ThermoFisher) at room temperature for 1h. Cells were washed thrice with PBS/0.1% Triton-X and imaged using an ECHO revolve 2 microscope. DAPI (ThermoFisher) was used as a nuclear stain.

##### *PSEN1*^G206A^ knock-in via CRISPR/Cas9

The G206A mutation was knocked into both alleles of *PSEN1* in IMR90 cl.4 iPSCs using CRISPR/Cas9 as has been done previously for the APP^Lon^ mutation.^86^ In brief, sgRNAs targeting exon 7 of *PSEN1* were designed (Deskgen.com) and subcloned into MLM3636 vector, a gift from Keith Joung (Addgene plasmid # 43860; http://n2t.net/addgene:43860; RRID:Addgene_43860). sgRNAs were tested for efficacy in HEK293 (ATCC) using a genomic cleavage kit (GeneArt, ThermoFisher). The best sgRNA was used for editing and had the sequence: TTCACTGGAAAGGTCCACTT. The ssDNA HDR template was designed with a silent CRISPR blocking mutation at the PAM site which added a de novo EcoR V restriction site in addition to the G206A mutation. Introduced changes are shown below in bolded under case in the HDR sequence. All oligos were for genome editing were ordered from Integrated DNA technologies. HDR: mutant sequence: GAAGTGTTTAAAACCTATAACGTTGCTGTGGACTACATTACTGTTGCACTCCTGATCTGGAATTT TG**c**TGTGGTGGGAATGAT**a**TC**g**ATTCACTGGAAAGGTCCACTTCGACTCCAGCAGGCATATC, eSpCas9(1.1), a gift from Feng Zhang^87^ (Addgene plasmid # 71814; http://n2t.net/addgene:71814; RRID:Addgene_71814), ssDNA HDR template, sgRNA and GFP plasmid (Lonza) were electroporated (Lonza nucleofector) into feeder-free iPSCs, followed by cell sorting on GFP signal and plating at low density on MEFs (MTI-GlobalStem). Individual clones were manually picked and transferred to 96-well format plate, which was split into duplicate plates and high and low density. gDNA was collected from high density plates as previously described.^88^ For each clone, exon 7 of *PSEN1* was amplified and screened by restriction digest by EcoR V (New England Biolabs). Sanger sequencing was used to confirm the mutation (Azenta), and successful clones were expanded and banked. *PSEN1*^G206A^ knock-in lines were analyzed and demonstrated to have a normal karyotype (Cell Line Genetics) and to be bona fide homozygous knock-ins rather hemizygous by a quantitative genotyping PCR method.^68^ Cl.95 was used in the present study.

##### RNA-seq analysis of neurons

RNA from control hiPSC-neurons treated with either control or siRNA targeting *SH3RF3* were sent for bulk RNA-seq by Azenta as has been done previously^89^. We compared the differential gene expression levels in siRNA vs. controls as implemented in the NetworkAnalyst (https://www.networkanalyst.ca), a web-based comprehensive gene expression profiling for the two groups.

##### Statistics

All experiments were repeated independently and information about number of repeated experiments, replicates, error bars, sample size the statistical test used, and measure of significance are provided in corresponding figure legends. Statistical analyses were performed in Prism software version 9.0 (Graphpad). Unpaired t-test was used for comparing the means of two groups. One-way ANOVA was used for experiments with single independent variable and two-way ANOVA was used for comparing the means of more than two groups, followed Tukey’s post-hoc test.

## References

1. Selkoe, D.J., and Hardy, J. (2016). The amyloid hypothesis of Alzheimer’s disease at 25 years. EMBO Mol Med 8, 595–608. 10.15252/emmm.201606210.

2. van Dyck, C.H., Swanson, C.J., Aisen, P., Bateman, R.J., Chen, C., Gee, M., Kanekiyo, M., Li, D., Reyderman, L., Cohen, S., et al. (2023). Lecanemab in Early Alzheimer’s Disease. N Engl J Med 388, 9–21. 10.1056/NEJMoa2212948.

3. Soderberg, L., Johannesson, M., Nygren, P., Laudon, H., Eriksson, F., Osswald, G., Moller, C., and Lannfelt, L. (2023). Lecanemab, Aducanumab, and Gantenerumab - Binding Profiles to Different Forms of Amyloid-Beta Might Explain Efficacy and Side Effects in Clinical Trials for Alzheimer’s Disease. Neurotherapeutics 20, 195–206. 10.1007/s13311-022-01308-6.

4. Lee, J.H., Cheng, R., Vardarajan, B., Lantigua, R., Reyes-Dumeyer, D., Ortmann, W., Graham, R.R., Bhangale, T., Behrens, T.W., Medrano, M., et al. (2015). Genetic Modifiers of Age at Onset in Carriers of the G206A Mutation in PSEN1 With Familial Alzheimer Disease Among Caribbean Hispanics. JAMA Neurol 72, 1043–1051. 10.1001/jamaneurol.2015.1424.

5. Lopera, F., Marino, C., Chandrahas, A.S., O’Hare, M., Villalba-Moreno, N.D., Aguillon, D., Baena, A., Sanchez, J.S., Vila-Castelar, C., Ramirez Gomez, L., et al. (2023). Resilience to autosomal dominant Alzheimer’s disease in a Reelin-COLBOS heterozygous man. Nat Med 29, 1243–1252. 10.1038/s41591-023-02318-3.

6. Schwartz, M.L.B., Williams, M.S., and Murray, M.F. (2017). Adding Protective Genetic Variants to Clinical Reporting of Genomic Screening Results: Restoring Balance. JAMA 317, 1527–1528. 10.1001/jama.2017.1533.

7. Park, Y.P., and Kellis, M. (2021). CoCoA-diff: counterfactual inference for single-cell gene expression analysis. Genome Biol 22, 228. 10.1186/s13059-021-02438-4.

8. Jia, P., Zhao, Z., Hulgan, T., Bush, W.S., Samuels, D.C., Bloss, C.S., Heaton, R.K., Ellis, R.J., Schork, N., Marra, C.M., et al. (2017). Genome-wide association study of HIV-associated neurocognitive disorder (HAND): A CHARTER group study. Am J Med Genet B Neuropsychiatr Genet 174, 413–426. 10.1002/ajmg.b.32530.

9. Karkkainen, S., van der Linden, M., and Renkema, G.H. (2010). POSH2 is a RING finger E3 ligase with Rac1 binding activity through a partial CRIB domain. FEBS Lett 584, 3867–3872. 10.1016/j.febslet.2010.07.060.

10. Tapon, N., Nagata, K., Lamarche, N., and Hall, A. (1998). A new rac target POSH is an SH3-containing scaffold protein involved in the JNK and NF-kappaB signalling pathways. EMBO J 17, 1395–1404. 10.1093/emboj/17.5.1395.

11. Zhang, P., Liu, Y., Lian, C., Cao, X., Wang, Y., Li, X., Cong, M., Tian, P., Zhang, X., Wei, G., et al. (2020). SH3RF3 promotes breast cancer stem-like properties via JNK activation and PTX3 upregulation. Nat Commun 11, 2487. 10.1038/s41467-020-16051-9.

12. Xu, Z., Kukekov, N.V., and Greene, L.A. (2003). POSH acts as a scaffold for a multiprotein complex that mediates JNK activation in apoptosis. EMBO J 22, 252–261. 10.1093/emboj/cdg021.

13. Kukekov, N.V., Xu, Z., and Greene, L.A. (2006). Direct interaction of the molecular scaffolds POSH and JIP is required for apoptotic activation of JNKs. J Biol Chem 281, 15517–15524. 10.1074/jbc.M601056200.

14. Wilhelm, M., Kukekov, N.V., Schmit, T.L., Biagas, K.V., Sproul, A.A., Gire, S., Maes, M.E., Xu, Z., and Greene, L.A. (2012). Sh3rf2/POSHER protein promotes cell survival by ring-mediated proteasomal degradation of the c-Jun N-terminal kinase scaffold POSH (Plenty of SH3s) protein. J Biol Chem 287, 2247–2256. 10.1074/jbc.M111.269431.

15. Kim, E.K., and Choi, E.J. (2010). Pathological roles of MAPK signaling pathways in human diseases. Biochim Biophys Acta 1802, 396–405. 10.1016/j.bbadis.2009.12.009.

16. Yarza, R., Vela, S., Solas, M., and Ramirez, M.J. (2015). c-Jun N-terminal Kinase (JNK) Signaling as a Therapeutic Target for Alzheimer’s Disease. Front Pharmacol 6, 321. 10.3389/fphar.2015.00321.

17. Sun, E., Motolani, A., Campos, L., and Lu, T. (2022). The Pivotal Role of NF-kB in the Pathogenesis and Therapeutics of Alzheimer’s Disease. Int J Mol Sci 23. 10.3390/ijms23168972.

18. Shi, Z.M., Han, Y.W., Han, X.H., Zhang, K., Chang, Y.N., Hu, Z.M., Qi, H.X., Ting, C., Zhen, Z., and Hong, W. (2016). Upstream regulators and downstream effectors of NF-kappaB in Alzheimer’s disease. J Neurol Sci 366, 127–134. 10.1016/j.jns.2016.05.022.

19. Lee, M.S., Kao, S.C., Lemere, C.A., Xia, W., Tseng, H.C., Zhou, Y., Neve, R., Ahlijanian, M.K., and Tsai, L.H. (2003). APP processing is regulated by cytoplasmic phosphorylation. J Cell Biol 163, 83–95. 10.1083/jcb.200301115.

20. Colombo, A., Bastone, A., Ploia, C., Sclip, A., Salmona, M., Forloni, G., and Borsello, T. (2009). JNK regulates APP cleavage and degradation in a model of Alzheimer’s disease. Neurobiol Dis 33, 518–525. 10.1016/j.nbd.2008.12.014.

21. Yoon, S.O., Park, D.J., Ryu, J.C., Ozer, H.G., Tep, C., Shin, Y.J., Lim, T.H., Pastorino, L., Kunwar, A.J., Walton, J.C., et al. (2012). JNK3 perpetuates metabolic stress induced by Abeta peptides. Neuron 75, 824–837. 10.1016/j.neuron.2012.06.024.

22. Yoshida, H., Hastie, C.J., McLauchlan, H., Cohen, P., and Goedert, M. (2004). Phosphorylation of microtubule-associated protein tau by isoforms of c-Jun N-terminal kinase (JNK). J Neurochem 90, 352–358. 10.1111/j.1471-4159.2004.02479.x.

23. Reynolds, C.H., Utton, M.A., Gibb, G.M., Yates, A., and Anderton, B.H. (1997). Stress-activated protein kinase/c-jun N-terminal kinase phosphorylates tau protein. J Neurochem 68, 1736–1744. 10.1046/j.1471-4159.1997.68041736.x.

24. Tamagno, E., Guglielmotto, M., Giliberto, L., Vitali, A., Borghi, R., Autelli, R., Danni, O., and Tabaton, M. (2009). JNK and ERK1/2 pathways have a dual opposite effect on the expression of BACE1. Neurobiol Aging 30, 1563–1573. 10.1016/j.neurobiolaging.2007.12.015.

25. Troy, C.M., Rabacchi, S.A., Xu, Z., Maroney, A.C., Connors, T.J., Shelanski, M.L., and Greene, L.A. (2001). beta-Amyloid-induced neuronal apoptosis requires c-Jun N-terminal kinase activation. J Neurochem 77, 157–164. 10.1046/j.1471-4159.2001.t01-1-00218.x.

26. Dong, W., Embury, C.M., Lu, Y., Whitmire, S.M., Dyavarshetty, B., Gelbard, H.A., Gendelman, H.E., and Kiyota, T. (2016). The mixed-lineage kinase 3 inhibitor URMC-099 facilitates microglial amyloid-beta degradation. J Neuroinflammation 13, 184. 10.1186/s12974-016-0646-z.

27. Zhong, L., Zhang, Z.L., Li, X., Liao, C., Mou, P., Wang, T., Wang, Z., Wang, Z., Wei, M., Xu, H., et al. (2017). TREM2/DAP12 Complex Regulates Inflammatory Responses in Microglia via the JNK Signaling Pathway. Front Aging Neurosci 9, 204. 10.3389/fnagi.2017.00204.

28. Ruganzu, J.B., Peng, X., He, Y., Wu, X., Zheng, Q., Ding, B., Lin, C., Guo, H., Yang, Z., Zhang, X., and Yang, W. (2022). Downregulation of TREM2 expression exacerbates neuroinflammatory responses through TLR4-mediated MAPK signaling pathway in a transgenic mouse model of Alzheimer’s disease. Mol Immunol 142, 22–36. 10.1016/j.molimm.2021.12.018.

29. Chen, C.H., Zhou, W., Liu, S., Deng, Y., Cai, F., Tone, M., Tone, Y., Tong, Y., and Song, W. (2012). Increased NF-kappaB signalling up-regulates BACE1 expression and its therapeutic potential in Alzheimer’s disease. Int J Neuropsychopharmacol 15, 77–90. 10.1017/S1461145711000149.

30. Mancuso, R., Fattorelli, N., Martinez-Muriana, A., Davis, E., Wolfs, L., Van Den Daele, J., Geric, I., Premereur, J., Polanco, P., Bijnens, B., et al. (2024). Xenografted human microglia display diverse transcriptomic states in response to Alzheimer’s disease-related amyloid-beta pathology. Nat Neurosci 27, 886–900. 10.1038/s41593-024-01600-y.

31. Lee, J.H., Mayeux, R., Mayo, D., Mo, J., Santana, V., Williamson, J., Flaquer, A., Ciappa, A., Rondon, H., Estevez, P., et al. (2004). Fine mapping of 10q and 18q for familial Alzheimer’s disease in Caribbean Hispanics. Mol Psychiatry 9, 1042–1051. 10.1038/sj.mp.4001538.

32. Tang, M.X., Stern, Y., Marder, K., Bell, K., Gurland, B., Lantigua, R., Andrews, H., Feng, L., Tycko, B., and Mayeux, R. (1998). The APOE-epsilon4 allele and the risk of Alzheimer disease among African Americans, whites, and Hispanics. JAMA 279, 751–755. 10.1001/jama.279.10.751.

33. Song, S., Ashok, A., Williams, D., Kaufman, M., Duff, K., and Sproul, A. (2021). Efficient Derivation of Excitatory and Inhibitory Neurons from Human Pluripotent Stem Cells Stably Expressing Direct Reprogramming Factors. Curr Protoc 1, e141. 10.1002/cpz1.141.

34. Huang, Y.A., Zhou, B., Wernig, M., and Sudhof, T.C. (2017). ApoE2, ApoE3, and ApoE4 Differentially Stimulate APP Transcription and Abeta Secretion. Cell 168, 427–441 e421. 10.1016/j.cell.2016.12.044.

35. Vogel, J., Anand, V.S., Ludwig, B., Nawoschik, S., Dunlop, J., and Braithwaite, S.P. (2009). The JNK pathway amplifies and drives subcellular changes in tau phosphorylation. Neuropharmacology 57, 539–550. 10.1016/j.neuropharm.2009.07.021.

36. Kim, D., Su, J., and Cotman, C.W. (1999). Sequence of neurodegeneration and accumulation of phosphorylated tau in cultured neurons after okadaic acid treatment. Brain Res 839, 253–262. 10.1016/s0006-8993(99)01724-2.

37. Hasegawa, M., Jakes, R., Crowther, R.A., Lee, V.M., Ihara, Y., and Goedert, M. (1996). Characterization of mAb AP422, a novel phosphorylation-dependent monoclonal antibody against tau protein. FEBS Lett 384, 25–30. 10.1016/0014-5793(96)00271-2.

38. Deters, N., Ittner, L.M., and Gotz, J. (2008). Divergent phosphorylation pattern of tau in P301L tau transgenic mice. Eur J Neurosci 28, 137–147. 10.1111/j.1460-9568.2008.06318.x.

39. Haase, C., Stieler, J.T., Arendt, T., and Holzer, M. (2004). Pseudophosphorylation of tau protein alters its ability for self-aggregation. J Neurochem 88, 1509–1520. 10.1046/j.1471-4159.2003.02287.x.

40. Montoliu-Gaya, L., Benedet, A.L., Tissot, C., Vrillon, A., Ashton, N.J., Brum, W.S., Lantero-Rodriguez, J., Stevenson, J., Nilsson, J., Sauer, M., et al. (2023). Mass spectrometric simultaneous quantification of tau species in plasma shows differential associations with amyloid and tau pathologies. Nat Aging 3, 661–669. 10.1038/s43587-023-00405-1.

41. Barthelemy, N.R., Mallipeddi, N., Moiseyev, P., Sato, C., and Bateman, R.J. (2019). Tau Phosphorylation Rates Measured by Mass Spectrometry Differ in the Intracellular Brain vs. Extracellular Cerebrospinal Fluid Compartments and Are Differentially Affected by Alzheimer’s Disease. Front Aging Neurosci 11, 121. 10.3389/fnagi.2019.00121.

42. Huang da, W., Sherman, B.T., and Lempicki, R.A. (2009). Systematic and integrative analysis of large gene lists using DAVID bioinformatics resources. Nat Protoc 4, 44–57. 10.1038/nprot.2008.211.

43. Merighi, S., Nigro, M., Travagli, A., and Gessi, S. (2022). Microglia and Alzheimer’s Disease. Int J Mol Sci 23. 10.3390/ijms232112990.

44. Mehan, S., Meena, H., Sharma, D., and Sankhla, R. (2011). JNK: a stress-activated protein kinase therapeutic strategies and involvement in Alzheimer’s and various neurodegenerative abnormalities. J Mol Neurosci 43, 376–390. 10.1007/s12031-010-9454-6.

45. Zhang, Y., Sloan, S.A., Clarke, L.E., Caneda, C., Plaza, C.A., Blumenthal, P.D., Vogel, H., Steinberg, G.K., Edwards, M.S., Li, G., et al. (2016). Purification and Characterization of Progenitor and Mature Human Astrocytes Reveals Transcriptional and Functional Differences with Mouse. Neuron 89, 37–53. 10.1016/j.neuron.2015.11.013.

46. Zhang, Y., Chen, K., Sloan, S.A., Bennett, M.L., Scholze, A.R., O’Keeffe, S., Phatnani, H.P., Guarnieri, P., Caneda, C., Ruderisch, N., et al. (2014). An RNA-sequencing transcriptome and splicing database of glia, neurons, and vascular cells of the cerebral cortex. J Neurosci 34, 11929–11947. 10.1523/JNEUROSCI.1860-14.2014.

47. Levine, K.S., Leonard, H.L., Blauwendraat, C., Iwaki, H., Johnson, N., Bandres-Ciga, S., Ferrucci, L., Faghri, F., Singleton, A.B., and Nalls, M.A. (2023). Virus exposure and neurodegenerative disease risk across national biobanks. Neuron 111, 1086–1093 e1082. 10.1016/j.neuron.2022.12.029.

48. Steel, A.J., and Eslick, G.D. (2015). Herpes Viruses Increase the Risk of Alzheimer’s Disease: A Meta-Analysis. J Alzheimers Dis 47, 351–364. 10.3233/JAD-140822.

49. Liu, N., Jiang, X., and Li, H. (2023). The viral hypothesis in Alzheimer’s disease: SARS-CoV-2 on the cusp. Front Aging Neurosci 15, 1129640. 10.3389/fnagi.2023.1129640.

50. Priemer, D.S., Rhodes, C.H., Karlovich, E., Perl, D.P., and Goldman, J.E. (2022). Abeta Deposits in the Neocortex of Adult and Infant Hypoxic Brains, Including in Cases of COVID-19. J Neuropathol Exp Neurol 81, 988–995. 10.1093/jnen/nlac095.

51. Shen, W.B., Elahi, M., Logue, J., Yang, P., Baracco, L., Reece, E.A., Wang, B., Li, L., Blanchard, T.G., Han, Z., et al. (2022). SARS-CoV-2 invades cognitive centers of the brain and induces Alzheimer’s-like neuropathology. bioRxiv. 10.1101/2022.01.31.478476.

52. Kumar, V. (2019). Toll-like receptors in the pathogenesis of neuroinflammation. J Neuroimmunol 332, 16–30. 10.1016/j.jneuroim.2019.03.012.

53. Wang, W.Y., Tan, M.S., Yu, J.T., and Tan, L. (2015). Role of pro-inflammatory cytokines released from microglia in Alzheimer’s disease. Ann Transl Med 3, 136. 10.3978/j.issn.2305-5839.2015.03.49.

54. <Oligodendrocytes damage in Alzheimer’s disease Beta amyloid toxicity and inflammation.pdf>.

55. Cai, Y., Liu, J., Wang, B., Sun, M., and Yang, H. (2022). Microglia in the Neuroinflammatory Pathogenesis of Alzheimer’s Disease and Related Therapeutic Targets. Front Immunol 13, 856376. 10.3389/fimmu.2022.856376.

56. He, Y., Taylor, N., Yao, X., and Bhattacharya, A. (2021). Mouse primary microglia respond differently to LPS and poly(I:C) in vitro. Sci Rep 11, 10447. 10.1038/s41598-021-89777-1.

57. McQuade, A., Coburn, M., Tu, C.H., Hasselmann, J., Davtyan, H., and Blurton-Jones, M. (2018). Development and validation of a simplified method to generate human microglia from pluripotent stem cells. Mol Neurodegener 13, 67. 10.1186/s13024-018-0297-x.

58. Maroney, A.C., Finn, J.P., Connors, T.J., Durkin, J.T., Angeles, T., Gessner, G., Xu, Z., Meyer, S.L., Savage, M.J., Greene, L.A., et al. (2001). Cep-1347 (KT7515), a semisynthetic inhibitor of the mixed lineage kinase family. J Biol Chem 276, 25302–25308. 10.1074/jbc.M011601200.

59. Parkinson Study Group, P.I. (2007). Mixed lineage kinase inhibitor CEP-1347 fails to delay disability in early Parkinson disease. Neurology 69, 1480–1490. 10.1212/01.wnl.0000277648.63931.c0.

60. Waelchli, R., Bollbuck, B., Bruns, C., Buhl, T., Eder, J., Feifel, R., Hersperger, R., Janser, P., Revesz, L., Zerwes, H.G., and Schlapbach, A. (2006). Design and preparation of 2-benzamido-pyrimidines as inhibitors of IKK. Bioorg Med Chem Lett 16, 108–112. 10.1016/j.bmcl.2005.09.035.

61. Zhang, Q.G., Wang, R., Han, D., Dong, Y., and Brann, D.W. (2009). Role of Rac1 GTPase in JNK signaling and delayed neuronal cell death following global cerebral ischemia. Brain Res 1265, 138–147. 10.1016/j.brainres.2009.01.033.

62. <Discovery and characterization of small molecule Rac1 inhibitors.pdf>.

63. Dhawan, G., Floden, A.M., and Combs, C.K. (2012). Amyloid-beta oligomers stimulate microglia through a tyrosine kinase dependent mechanism. Neurobiol Aging 33, 2247–2261. 10.1016/j.neurobiolaging.2011.10.027.

64. Parajuli, B., Sonobe, Y., Horiuchi, H., Takeuchi, H., Mizuno, T., and Suzumura, A. (2013). Oligomeric amyloid beta induces IL-1beta processing via production of ROS: implication in Alzheimer’s disease. Cell Death Dis 4, e975. 10.1038/cddis.2013.503.

65. Fa, M., Orozco, I.J., Francis, Y.I., Saeed, F., Gong, Y., and Arancio, O. (2010). Preparation of oligomeric beta-amyloid 1-42 and induction of synaptic plasticity impairment on hippocampal slices. J Vis Exp. 10.3791/1884.

66. Quiroga, I.Y., Cruikshank, A.E., Bond, M.L., Reed, K.S.M., Evangelista, B.A., Tseng, J.H., Ragusa, J.V., Meeker, R.B., Won, H., Cohen, S., et al. (2022). Synthetic amyloid beta does not induce a robust transcriptional response in innate immune cell culture systems. J Neuroinflammation 19, 99. 10.1186/s12974-022-02459-1.

67. Konttinen, H., Cabral-da-Silva, M.E.C., Ohtonen, S., Wojciechowski, S., Shakirzyanova, A., Caligola, S., Giugno, R., Ishchenko, Y., Hernandez, D., Fazaludeen, M.F., et al. (2019). PSEN1DeltaE9, APPswe, and APOE4 Confer Disparate Phenotypes in Human iPSC-Derived Microglia. Stem Cell Reports 13, 669–683. 10.1016/j.stemcr.2019.08.004.

68. Weisheit, I., Kroeger, J.A., Malik, R., Klimmt, J., Crusius, D., Dannert, A., Dichgans, M., and Paquet, D. (2020). Detection of Deleterious On-Target Effects after HDR-Mediated CRISPR Editing. Cell Rep 31, 107689. 10.1016/j.celrep.2020.107689.

69. Ma, Q.L., Yang, F., Rosario, E.R., Ubeda, O.J., Beech, W., Gant, D.J., Chen, P.P., Hudspeth, B., Chen, C., Zhao, Y., et al. (2009). Beta-amyloid oligomers induce phosphorylation of tau and inactivation of insulin receptor substrate via c-Jun N-terminal kinase signaling: suppression by omega-3 fatty acids and curcumin. J Neurosci 29, 9078–9089. 10.1523/JNEUROSCI.1071-09.2009.

70. Guillozet-Bongaarts, A.L., Cahill, M.E., Cryns, V.L., Reynolds, M.R., Berry, R.W., and Binder, L.I. (2006). Pseudophosphorylation of tau at serine 422 inhibits caspase cleavage: in vitro evidence and implications for tangle formation in vivo. J Neurochem 97, 1005–1014. 10.1111/j.1471-4159.2006.03784.x.

71. Wunderlich, P., Glebov, K., Kemmerling, N., Tien, N.T., Neumann, H., and Walter, J. (2013). Sequential proteolytic processing of the triggering receptor expressed on myeloid cells-2 (TREM2) protein by ectodomain shedding and gamma-secretase-dependent intramembranous cleavage. J Biol Chem 288, 33027–33036. 10.1074/jbc.M113.517540.

72. Salminen, A., Kauppinen, A., and Kaarniranta, K. (2017). Hypoxia/ischemia activate processing of Amyloid Precursor Protein: impact of vascular dysfunction in the pathogenesis of Alzheimer’s disease. J Neurochem 140, 536–549. 10.1111/jnc.13932.

73. Dolan, M.J., Therrien, M., Jereb, S., Kamath, T., Gazestani, V., Atkeson, T., Marsh, S.E., Goeva, A., Lojek, N.M., Murphy, S., et al. (2023). Exposure of iPSC-derived human microglia to brain substrates enables the generation and manipulation of diverse transcriptional states in vitro. Nat Immunol 24, 1382–1390. 10.1038/s41590-023-01558-2.

74. Lund, S., Porzgen, P., Mortensen, A.L., Hasseldam, H., Bozyczko-Coyne, D., Morath, S., Hartung, T., Bianchi, M., Ghezzi, P., Bsibsi, M., et al. (2005). Inhibition of microglial inflammation by the MLK inhibitor CEP-1347. J Neurochem 92, 1439–1451. 10.1111/j.1471-4159.2005.03014.x.

75. Kiyota, T., Machhi, J., Lu, Y., Dyavarshetty, B., Nemati, M., Zhang, G., Mosley, R.L., Gelbard, H.A., and Gendelman, H.E. (2018). URMC-099 facilitates amyloid-beta clearance in a murine model of Alzheimer’s disease. J Neuroinflammation 15, 137. 10.1186/s12974-018-1172-y.

76. Maroney, A.C., Finn, J.P., Bozyczko-Coyne, D., O’Kane, T.M., Neff, N.T., Tolkovsky, A.M., Park, D.S., Yan, C.Y., Troy, C.M., and Greene, L.A. (1999). CEP-1347 (KT7515), an inhibitor of JNK activation, rescues sympathetic neurons and neuronally differentiated PC12 cells from death evoked by three distinct insults. J Neurochem 73, 1901–1912.

77. Waldmeier, P., Bozyczko-Coyne, D., Williams, M., and Vaught, J.L. (2006). Recent clinical failures in Parkinson’s disease with apoptosis inhibitors underline the need for a paradigm shift in drug discovery for neurodegenerative diseases. Biochem Pharmacol 72, 1197–1206. 10.1016/j.bcp.2006.06.031.

78. Chen, X., Rzhetskaya, M., Kareva, T., Bland, R., During, M.J., Tank, A.W., Kholodilov, N., and Burke, R.E. (2008). Antiapoptotic and trophic effects of dominant-negative forms of dual leucine zipper kinase in dopamine neurons of the substantia nigra in vivo. J Neurosci 28, 672–680. 10.1523/JNEUROSCI.2132-07.2008.

79. Sariya, S., Lee, J.H., Mayeux, R., Vardarajan, B.N., Reyes-Dumeyer, D., Manly, J.J., Brickman, A.M., Lantigua, R., Medrano, M., Jimenez-Velazquez, I.Z., and Tosto, G. (2019). Rare Variants Imputation in Admixed Populations: Comparison Across Reference Panels and Bioinformatics Tools. Front Genet 10, 239. 10.3389/fgene.2019.00239.

80. Sun, L., Dimitromanolakis, A., and Chen, W.M. (2017). Identifying Cryptic Relationships. Methods Mol Biol 1666, 45–60. 10.1007/978-1-4939-7274-6_4.

81. Yu, J., Vodyanik, M.A., Smuga-Otto, K., Antosiewicz-Bourget, J., Frane, J.L., Tian, S., Nie, J., Jonsdottir, G.A., Ruotti, V., Stewart, R., et al. (2007). Induced pluripotent stem cell lines derived from human somatic cells. Science 318, 1917–1920. 10.1126/science.1151526.

82. Abud, E.M., Ramirez, R.N., Martinez, E.S., Healy, L.M., Nguyen, C.H.H., Newman, S.A., Yeromin, A.V., Scarfone, V.M., Marsh, S.E., Fimbres, C., et al. (2017). iPSC-Derived Human Microglia-like Cells to Study Neurological Diseases. Neuron 94, 278–293 e279. 10.1016/j.neuron.2017.03.042.

83. Lafaille, F.G., Pessach, I.M., Zhang, S.Y., Ciancanelli, M.J., Herman, M., Abhyankar, A., Ying, S.W., Keros, S., Goldstein, P.A., Mostoslavsky, G., et al. (2012). Impaired intrinsic immunity to HSV-1 in human iPSC-derived TLR3-deficient CNS cells. Nature 491, 769–773. 10.1038/nature11583.

84. Kim, Y.A., Siddiqui, T., Blaze, J., Cosacak, M.I., Winters, T., Kumar, A., Tein, E., Sproul, A.A., Teich, A.F., Bartolini, F., et al. (2023). RNA methyltransferase NSun2 deficiency promotes neurodegeneration through epitranscriptomic regulation of tau phosphorylation. Acta Neuropathol 145, 29–48. 10.1007/s00401-022-02511-7.

85. Stine, W.B., Jr., Dahlgren, K.N., Krafft, G.A., and LaDu, M.J. (2003). In vitro characterization of conditions for amyloid-beta peptide oligomerization and fibrillogenesis. J Biol Chem 278, 11612–11622. 10.1074/jbc.M210207200.

86. Sun, J., Carlson-Stevermer, J., Das, U., Shen, M., Delenclos, M., Snead, A.M., Koo, S.Y., Wang, L., Qiao, D., Loi, J., et al. (2019). CRISPR/Cas9 editing of APP C-terminus attenuates beta-cleavage and promotes alpha-cleavage. Nat Commun 10, 53. 10.1038/s41467-018-07971-8.

87. Slaymaker, I.M., Gao, L., Zetsche, B., Scott, D.A., Yan, W.X., and Zhang, F. (2016). Rationally engineered Cas9 nucleases with improved specificity. Science 351, 84–88. 10.1126/science.aad5227.

88. Paquet, D., Kwart, D., Chen, A., Sproul, A., Jacob, S., Teo, S., Olsen, K.M., Gregg, A., Noggle, S., and Tessier-Lavigne, M. (2016). Efficient introduction of specific homozygous and heterozygous mutations using CRISPR/Cas9. Nature 533, 125–129. 10.1038/nature17664.

89. Bartosch, A.M.W., Youth, E.H.H., Hansen, S., Wu, Y., Buchanan, H.M., Kaufman, M.E., Xiao, H., Koo, S.Y., Ashok, A., Sivakumar, S., et al. (2024). ZCCHC17 Modulates Neuronal RNA Splicing and Supports Cognitive Resilience in Alzheimer’s Disease. J Neurosci 44. 10.1523/JNEUROSCI.2324-22.2023.

