## Supplemental Table S1 for "Reduced *SH3RF3* may protect against Alzheimer’s disease by lowering microglial pro-inflammatory responses via modulation of JNK and NFkB signaling"

| **Supplement Table 1 related to Table 1. Replications: Demographic and clinical characteristics of of LOAD data, 4060 EFIGAs and 2058 WHICAPs** | | | | |
| --- | --- | --- | --- | --- |
| **Characteristics** | **Genotyped** | | | |
|  | **EFIGA** | | **WHICAP** | |
|  | **n** | **mean/%** | **n** | **mean/%** |
| Number of families | 434 |  |  |  |
| Number of family members | 2,114 |  |  |  |
| Singletons | 1,946 |  | 2,058 |  |
| Affection status |  |  |  |  |
| Affected | 1,913 | 47.1% | 1,033 | 50.2% |
| Unaffected | 2,147 | 52.9% | 1,025 | 49.8% |
| Proportion of females | 2,699 | 66.5% | 1,425 | 69.2% |
| Age |  |  |  |  |
| Age at onset age (affecteds) |  | 74.5(9.2) |  | 80.8(6.4) |
| Age at last examination (Unaffecteds) |  | 66.3(10.1) |  | 79.0(6.7) |
| APOE frequency |  |  |  |  |
| E4 | 1,996 | 24.6% | 544 | 13.2% |
| E3 | 5,712 | 70.3% | 3,264 | 79.3% |
| E2 | 412 | 5.1% | 308 | 7.5% |
| Mean years of education |  | 6.7(5.6) |  | 6.8(4.4) |
| Affecteds |  | 4.4(4.4) |  | 5.6(4.0) |
| Unaffecteds |  | 8.8(5.7) |  | 8.0(4.5) |
