## Supplemental Figure S2 for "Reduced *SH3RF3* may protect against Alzheimer’s disease by lowering microglial pro-inflammatory responses via modulation of JNK and NFkB signaling"

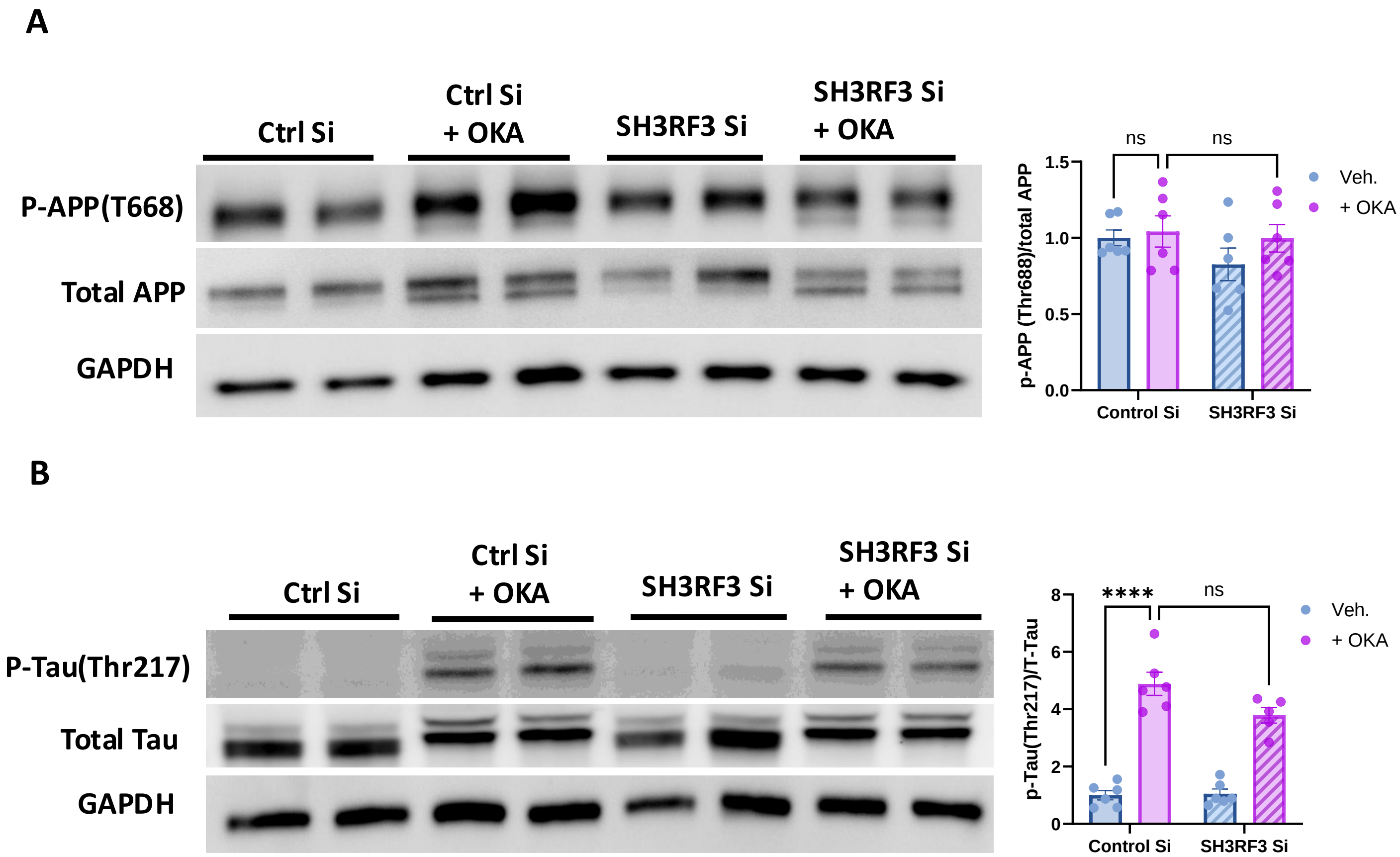


**Figure S2: SH3RF3 KD does not affect APP T668 phosphorylation and only trends to reduce p-Tau T217, related to Figure 2.** (A) Western blot analysis of phospho-APP (Thr668) phosphorylation with 3-hour OKA treatment with non-targeting siRNA and SH3RF3 siRNA in neurons. Representative blots and quantification is presented. Data is presented as mean±SEM with N=3 from two independent experiments, Comparison of control Si with OA and SH3RF3 Si with OA by two-way ANOVA. (B) Western blot analysis of Tau (Thr217) phosphorylation with 3-hour OKA treatment with non-targeting siRNA and SH3RF3 siRNA in neurons. Representative blots and quantification is presented. Data is presented as mean±SEM with N=3 from two independent experiments, Comparison of control Si with OA and SH3RF3 Si with OA by two-way ANOVA, ****<0.0001, NS
