## Supplemental Figure S3 for "Reduced *SH3RF3* may protect against Alzheimer’s disease by lowering microglial pro-inflammatory responses via modulation of JNK and NFkB signaling"

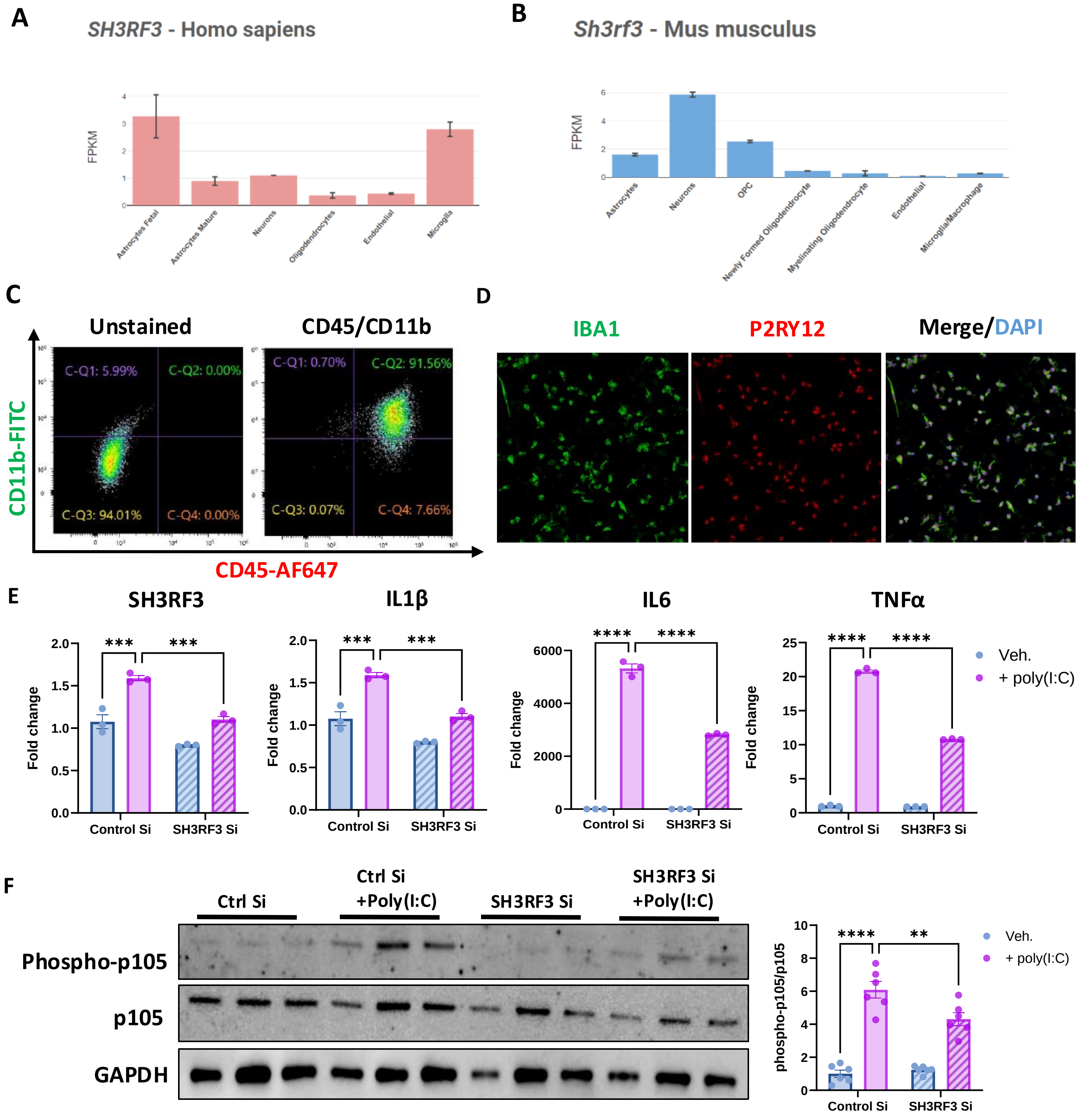


**Figure S3: iMGLs expressing microglial markers induces transcription of inflammatory cytokines and NFκB pathway activity with poly(I:C) treatment which is modulated by SH3RF3 knockdown.** (A-B) *SH3RF3* expression in CNS cells types determined by scRNAseq from published database (brainrnaseq.org) for human (A) and mice (B). (C) Characterization of iMGLs with flow cytometry showing cell stained for microglial markers CD45 and CD11b, CD45$^+$/CD11b$^+$ double positive cells are compared with unstained control. (D) Immunostaining for microglial markers IBA1 (green) and P2RY12 (red) for iMGLs, DAPI (blue) showed in merge is used as a nuclear stain (E) Representative experiment for qPCR (figure M1A) analysis of SH3RF3, IL1β, IL6 and TNFα with 2 hr poly(I:C) treatment with non-targeting siRNA and SH3RF3 siRNA in iMGLs, Data is presented as Mean±SEM with N=3 from one independent experiments, Comparison of control Si with poly(I:C) and SH3RF3 Si with poly(I:C) are made by ordinary two-way ANOVA, ns, ****<0.0001, ***<0.001 (F) Western blot analysis of phosphorylated p105 and total p105 with 6 hr poly(I:C) treatment with non-targeting siRNA and SH3RF3 siRNA in iMGLs. Representative blots and quantification is presented. Data is presented as Mean±SEM with N=3 from two independent experiments, Comparison of control Si with poly(I:C) and SH3RF3 Si with poly(I:C) by ordinary two-way ANOVA, ****<0.0001, **<0.01
