## Supplemental Figure S4 for "Reduced *SH3RF3* may protect against Alzheimer’s disease by lowering microglial pro-inflammatory responses via modulation of JNK and NFkB signaling"

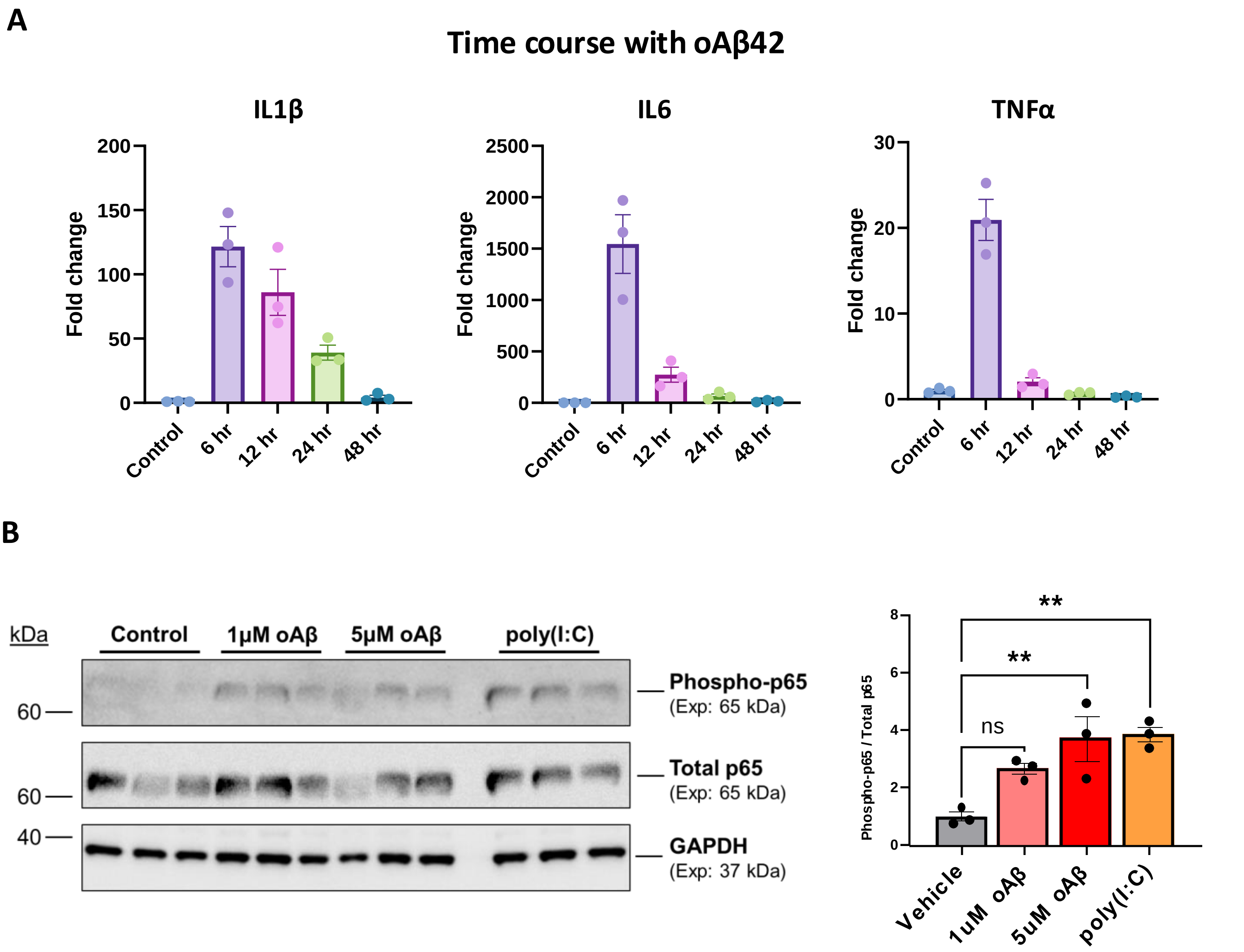


**Figure S4: Time course of cytokine transcription and NFκB pathway activation, related to Figure 4.** (A) qPCR analysis of IL1β, IL6 and TNFα with 5 uM oligomeric amyloid-β42 (oAβ) treatment in iMGLs at 6, 12, 24 and 48h compared with vehicle control (F12). Data is normalized to GAPDH expression and presented as mean±SEM with N=3. (B) Western blot analysis of phosphorylated and total p65 in iMGLs treated with vehicle (F12) 1 uM oAβ, 5 uM oAβ, or 25 ug/mL poly(I:C) for 3h. Data is presented as mean±SEM with N=3, Comparisons are made by ordinary one-way ANOVA, **<0.01, NS.
