## Supplemental Figure 5 for "Reduced *SH3RF3* may protect against Alzheimer’s disease by lowering microglial pro-inflammatory responses via modulation of JNK and NFkB signaling"

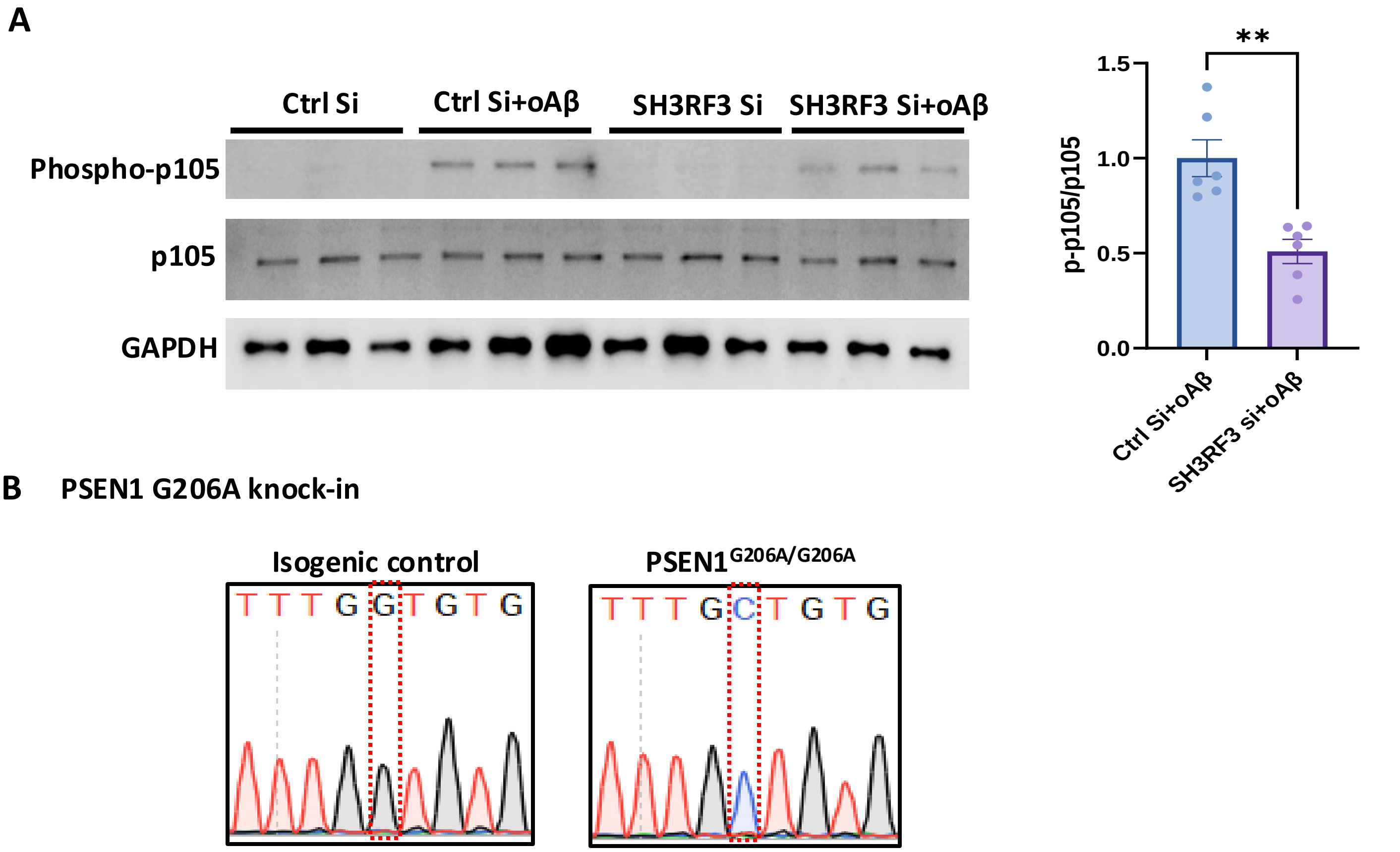


**Figure S5: SH3RF3 KD reduces phosphorylation of NFκB pathway component p105 induced by oβ42 treatment and CRISPR mediated knock-in of $PSEN1^{G206A}$, related to Figure 5.** (A) Western blot analysis of phosphorylated p105 and total p105 with 3h oAβ42 or vehicle treatment in presence of non-targeting siRNA or SH3RF3 siRNA in iMGLs. A representative blot and quantification for visible bands on oAβ42 conditions are shown. Data is presented as mean±SEM with N=3 from two independent experiments, Comparison of control Si with oAβ42 and SH3RF3 Si with oAβ42 by unpaired t-test, **<0.01. (B) Chromatographs from Sanger sequencing of exon 7 of PSEN1 show successful homozygous knock-in of the G206A mutation via CRISPR-mediated genome editing in the IMR90 (cl.4) hiPSC line. Red boxes marks site of the missense point mutation Chr14:73659420 G>C, which changes a glycine to an alanine.
